# A fluorescent non-hydrolyzable probe for the nucleotide binding sites of K_ATP_

**DOI:** 10.1101/2025.06.10.658839

**Authors:** Prisicilla Rubio, Shayla Q. Whitaker, Jon Ashby, Michael C. Puljung

## Abstract

Neuroendocrine ATP-sensitive K+ channels (K_ATP_) comprise four pore-forming subunits (Kir6.2), each associated with a modulatory sulfonylurea receptor subunit (SUR1). ATP/ADP binding to Kir6.2 inhibits K_ATP;_ MgATP/MgADP binding to two different sites on SUR1 promotes activation. As SUR1 is a member of the ABC transporter family of proteins, it can potentially hydrolyze MgATP to MgADP. Whether this activity is required for K_ATP_ activation remains controversial. Previous studies demonstrated that non-hydrolyzable ATP analogs do not activate K_ATP_, which may reflect an inability of these compounds to bind to SUR1, their inability to promote a conformational change in SUR1 that leads to channel activation, or a requirement for ATP hydrolysis during channel gating. To explore this further, we synthesized a fluorescent trinitrophenyl (TNP) derivative of the non-hydrolyzable ATP analog β,γ-methyleneadenosine 5’- triphosphate (AMP-PCP). Synthesis was verified by UV-visible absorbance, fluorescence spectroscopy, 1H nuclear magnetic resonance, and mass spectrometry. Purity was assessed by reversed-phase high-performance liquid chromatography. Real-time nucleotide binding to intact K_ATP_ channels in cell membranes was measured using FRET between channels labeled with a fluorescent, non-canonical amino acid and TNP-nucleotide derivatives. This technique provides sufficient spatial resolution to discriminate between binding to each site on K_ATP_. Using this approach we first established that TNP-ATP can bind to nucleotide binding site 1 on SUR1 in fluorescently labeled Kir6.2/SUR1 channels in unroofed membranes of HEK293T cells. We subsequently demonstrated that TNP-AMP-PCP binds to both nucleotide binding sites on SUR1 in the absence of Mg^2+^. AMP-PCP was able to compete with TNP-ATP for binding to NBS2, suggesting that it, too, binds NBS2. We conclude that the failure of non-hydrolyzable ATP analogs to activate K_ATP_ does not stem from an inability of these nucleotides to bind to the channel.

## INTRODUCTION

ATP sensitive K+ (K_ATP_) channels are endowed with a unique physiological gift: the ability to directly translate a change in a cell’s metabolic state into a change in its electrical excitability. Whereas K_ATP_ plays crucial roles in cardiovascular and neuronal tissues, its best understood task is in pancreatic β-cells, where changes in K_ATP_ open probability in response to an increase in blood sugar trigger insulin secretion (Puljung, 2023). Underlying this crucial function, mutations in the genes that encode the pancreatic K_ATP_ channel cause inherited diseases of insulin secretion including neonatal diabetes and congenital hyperinsulinism. Mutations that severely disrupt K_ATP_ function can also cause neuronal complications including developmental delay and epilepsy (Ashcroft et al., 2017).

The pancreatic K_ATP_ subtype is a hetero-octameric complex comprising four inward-rectifier K+ channel subunits (Kir6.2), each associated with a modulatory sulfonylurea receptor (SUR1) (Figure 1A,B) (Inagaki et al., 1997; Shyng and Nichols, 1997). Metabolic sensitivity is conferred on K_ATP_ through the binding of intracellular adenine nucleotides (ATP/ADP) to three classes of intracellular nucleotide-binding site (NBS), twelve sites in all, when accounting for the complex’s four-fold symmetry (Lee et al., 2017). ATP/ADP bind to four sites on Kir6.2 formed between adjacent subunits; nucleotide binding to Kir6.2 closes the channel in a reaction that does not require Mg^2+^ (Tucker et al., 1997). Nucleotides bind to two NBSs on SUR1, formed at the interface between its cytoplasmic nucleotide binding domains (NBDs, Figure 1C) (Lee et al., 2017). NBS1 is formed by the Walker motifs of NBD1 and the ABC signature sequence of NBD2; NBS2 is formed by the Walker motifs of NBD2 and the ABC signature sequence of NBD1. NBD dimerization drives a conformational change in the transmembrane domains of SUR1 that is coupled to an increase in K_ATP_’s open probability (Gribble et al., 1998). Whereas nucleotides bind to both NBSs of SUR1 in the presence or absence of Mg^2+^, Mg^2+^ is required to stabilize the NBD dimers and promote channel activation (Puljung et al., 2019).

**Figure 1.**
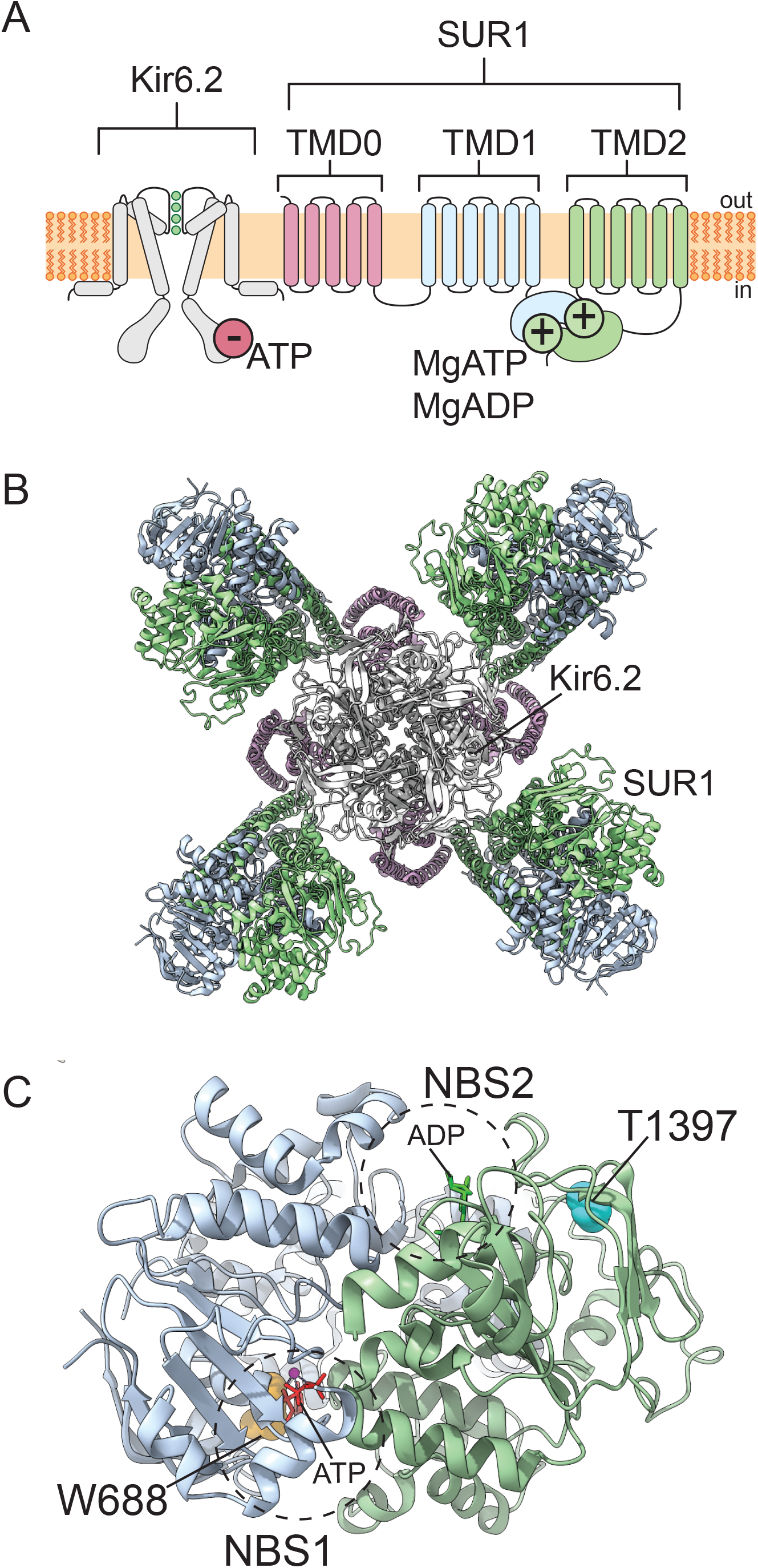
Structure of the pancreatic K_ATP_ channel. A. Cartoon showing the transmembrane topology of the pancreatic K_ATP_ channel. Two (of four) Kir6.2 subunits are shown along with one (of four) SUR1 subunits. Red and green circles mark the inhibitory and excitatory nucleotide binding sites (respectively). B. Cryo-EM structure of the pancreatic K_ATP_ channel determined in the presence of MgATP/ADP and viewed from the cytoplasmic side (Lee et al., 2017). The color scheme matches that of the schematic in panel A. PDB accession number 6C3O. C. The nucleotide binding domains of one SUR1 subunit viewed from the cytoplasmic side. ATP (bound to NBS1) and ADP (bound to NBS2) are shown as sticks. W688 and T1397 mark the sites that will be replaced with ANAP in our experiments. NBD1 is in blue and NBD2 is green.

SUR1 is a member of the ABC family of transporters (Aguilar-Bryan et al., 1995; Tusnády et al., 1997). Unlike most ABC proteins, SUR1 has not been shown to have any intrinsic transport activity, evolving only to modulate trafficking and opening of the Kir6.2 pore (Sakura et al., 1995; Inagaki et al., 1995). As a member of the ABC transporter family, SUR1 may have ATPase activity. NBS2, formed by the conserved, catalytic Walker_A_ and Walker_B_ motifs on NBD2 and the ABC signature sequence on NBD1 is competent to hydrolyze ATP. This is evident from *in vitro* ATPase activity in isolated NBD2 constructs and SUR1 subunits as well as a cryo-EM structure determined in the presence of MgATP, which surprisingly showed density for MgADP at NBS2 (Matsuo et al., 1999; Lee et al., 2017; de Wet et al., 2007). NBS1, on the other hand, is thought to be a degenerate site with no ATPase activity due to amino acid substitutions in its Walker_B_ motif and ABC signature sequence (Vedovato et al., 2015; Matsuo et al., 2000, 1999).

Degenerate sites are found in other ABC-family proteins including the closely related cystic fibrosis transmembrane conductance regulator (CFTR) channel and the bacterial transporter TM287/288 (Hohl et al., 2012; Aleksandrov et al., 2002).

Whereas SUR1 potentially retains the ability to hydrolyze ATP to ADP, the relevance of any such enzymatic activity for channel gating is unclear. K_ATP_ is directly activated by MgADP, implying that ATP hydrolysis is not required to provide a “power-stroke” coupled to channel activation (Proks et al., 2010). However, it remains an intriguing possibility that MgATP must first be hydrolyzed to MgADP to promote channel gating. Consistent with this idea, opening of cardiac K_ATP_ channels (formed by Kir6.2 and SUR2A) is promoted by vanadate, which stabilizes ABC proteins in a post-hydrolytic conformation (Zingman et al.). However, the time course reported for the current increase in vanadate is slow (taking 8.8 ± 0.9 min to develop) compared to channel gating, suggesting that hydrolysis may not be directly involved in channel opening.

For comparison, direct activation of Kir6.2-G334D/SUR1 by MgATP in inside-out patches was much faster, following a biexponential time course with time constants of 730 ms and 4.060 s (Proks et al., 2010). Further discounting the hypothesis that hydrolysis is required for gating, analysis of single-channel K_ATP_ currents do not show any out-of-equilibrium gating steps as one would expect for a process that involves irreversible hydrolysis (Choi et al., 2008; Csanady et al., 2010). Finally, ATP was able to promote a conformational change in isolated SUR1 subunits that displaced binding of tritiated glibenclamide (a K_ATP_ antagonist that binds SUR1) in the absence of Mg^2+^, i.e. under conditions that do not support hydrolysis (Ortiz et al., 2012).

However, this effect required very high ATP concentrations. Furthermore, these experiments were performed in isolated SUR1 subunits in *Pichia* membranes under equilibrium binding conditions, not in intact-functioning channels under physiological conditions.

Non-hydrolyzable ATP analogs can be used to probe the requirement for nucleotide hydrolysis by SUR1 in real-time using intact, functional channels in a membrane environment. Neither β,γ- methyleneadenosine 5’-triphosphate (AMP-PCP), which has a methylene group replacing the oxygen between the β and γ phosphates of ATP nor adenylyl-imidodiphosphate (AMP-PNP), which has an NH group at the same location, appear to support K_ATP_ activation (Dunne, 1989; Kozlowski and Ashford, 1992; Proks et al., 2010; Findlay, 1987; Ashcroft and Kakei, 1989; Treherne and Ashford, 1992; Jiang and Haddad, 1997; Schwanstecher and Panten, 1994; Schwanstecher et al., 1994; Hehl and Neumcke, 1994; Schwanstecher et al., 1992). Adenosine 5’-O-(3-thiotriphosphate) (ATP-γ-S) activates K_ATP_ channels for which the inhibitory nucleotide binding site on Kir6.2 has been removed (Kir6.2-G334D/SUR1) (Proks et al., 2010). However, this analog is weakly hydrolysable (Resetar and Chalovich, 1995).

There are at least three possible explanations for the inability of AMP-PCP and AMP-PNP to activate K_ATP_. 1) The nucleotides fail to bind to the NBSs of SUR1. 2) The nucleotides bind but fail to induce the conformational change (NBD dimerization) in SUR1 that promotes channel activation. 3). The nucleotides bind and dimerize the NBDs, but there is a *bona fide* requirement for ATP hydrolysis during the channel gating cycle. In this paper, we will address the first of these possibilities: binding. We have previously developed a technique that allows for the measurement of site-specific nucleotide binding to intact, functional K_ATP_ channels in the plasma membrane (Puljung et al., 2019; Usher et al., 2020b; a). This technique employs Förster resonance energy transfer (FRET) between the non-canonical amino acid L-3-(6- acetylnaphthalen-2-ylamino)-2-aminopropionic acid (ANAP), which can be incorporated directly into SUR1 using amber-stop-codon suppression, and fluorescent, trinitrophenyl (TNP) derivatives of ATP and ADP (Chatterjee et al., 2013; Puljung, 2021; Puljung et al., 2019; Usher et al., 2020a). The steep distance dependence of FRET allows for the discrimination of binding to each NBS. To evaluate non-hydrolyzable nucleotide effects on functional K_ATP_ channels in the plasma membrane, we synthesized, purified, and characterized a trinitrophenyl derivative of AMP-PCP (TNP-AMP-PCP). This derivative binds both NBSs of SUR1. Furthermore, we used a competition assay to demonstrate that non-fluorescent AMP-PCP can bind directly to NBS2. We conclude that the inability of non-hydrolyzable nucleotides to activate K_ATP_ does not arise from an inability to bind to SUR1 and either reflects an inability of these derivatives to change the conformation of SUR1 or a requirement for ATP hydrolysis to ADP to promote channel activation.

## MATERIALS AND METHODS

### TNP-AMP-PCP synthesis

TNP-AMP-PCP was synthesized using a modified version of the protocol for TNP-ATP synthesis published by Hiratsuka and Uchida (Figure 2A) (Hiratsuka and Uchida, 1973). 25 mg of the Na+ salt of β,γ-methyleneadenosine 5’-triphosphate (AMP-PCP; Enzo Biochem; Farmingdale, NY) were dissolved in 1 mL of deionized water and the concentration was verified by UV-vis spectroscopy (λ_max_ = 259 nm, ε = 15,400 M-1cm-1) using a Hitachi U-3010 UV-visible (Hitachi, Tokyo, Japan) and UV Solutions software. The measured concentration of AMP-PCP was 0.032 M (a total of 32 μmoles of AMP-PCP). 130 μL of a 1 M stock solution of 2,4,6-trinitrobenzenesulfonic acid (TNBS; Sigma-Aldrich, St. Louis, MO) in water were added to the AMP-PCP solution while stirring. The pH was adjusted to ∼9.5 with 240 μL of 1 M LiOH (Sigma-Aldrich). The reaction was stirred in the dark for four days at room temperature. Basic pH was maintained with daily additions of 1 M LiOH (an additional volume of 260 μL). Reaction progress was evident by the appearance of a deep orange color, resulting from the formation of a Meisenheimer complex subsequent to trinitrophenylation of the ribose (Azegami and Iwai, 1975).

**Figure 2.**
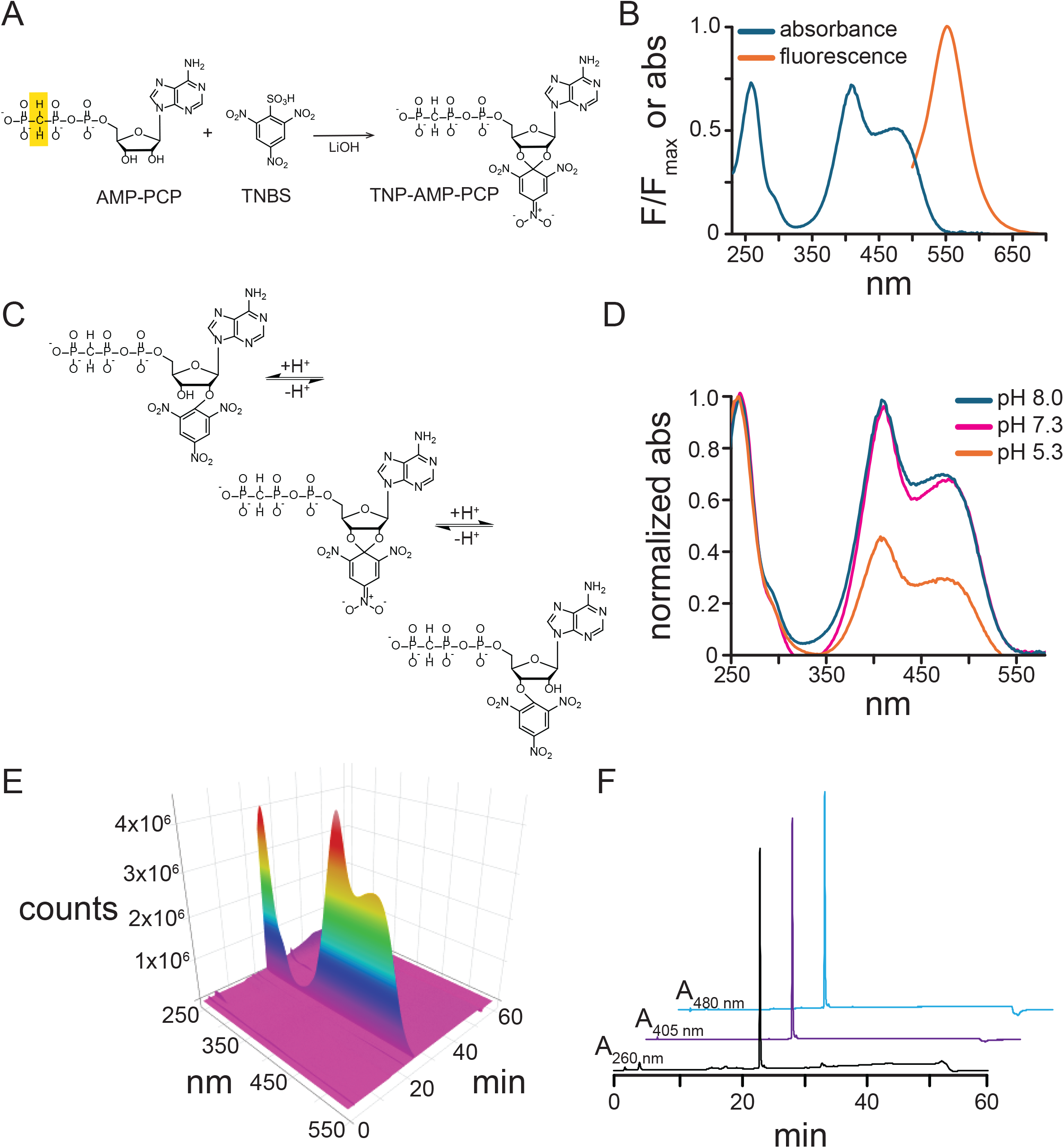
Synthesis and characterization of TNP-AMP-PCP. A. General reaction scheme for synthesis of TNP-AMP-PCP. The methylene group between the β and γ phosphates of AMP- PCP is highlighted in yellow. B. UV/Vis absorbance and normalized fluorescence emission spectra of purified TNP-AMP-PCP in 1M Tris, pH 8.0. C. Schematic depicting reversible protonation of the fluorescent Meisenheimer complex of TNP-AMP-PCP. D. UV/Vis spectrum of 42.15 µM TNP-AMP-PCP in Tris buffer (pH 8.0), HEPES buffer (pH 7.3), and MES buffer (pH 5.3). Data were normalized to the peak at 259 nm. E. Reversed-phase HPLC on purified TNP- AMP-PCP. The sample was applied to a C_18_ in 0.1 M TEAB buffer and eluted on a gradient of 0% to 60% acetonitrile in TEAB buffer. F. Cross sections of the chromatogram in C showing the elution of a single peak detected at 260 nm, 405 nm, and 480 nm. The traces are offset vertically and horizontally (by 5 min and 10 min for the 405 nm and 480 nm traces, respectively) for display purposes.

The product was purified by precipitation with 10 mL of 95% ethanol and 100 μL of 3 M sodium acetate, pH 5.2 (EMD Millipore Co.; Billerica, MA) and stored at -20 °C. The precipitate was collected by centrifugation at 13,500 x*g* in a Labnet Prism microcentrifuge (Labnet International; Edison, NJ). The product was washed twice with 95% ethanol, dried, and resuspended in deionized water for subsequent applications. Based on A259 (see below) the yield was estimated to be 79.3%.

We also attempted to synthesize trinitrophenyl derivatives of adenylyl-imidodiphosphate (AMP-PNP) and adenosine 5’-O-(3-thiotriphosphate) (ATP-γ-S). Whereas the reaction clearly proceeded and led to the formation of a Meisenheimer complex, subsequent mass spectrometric and thin-layer chromatography analyses suggested that one or more phosphate groups were lost during the reaction or purification process. Thus, these compounds were not pursued any further.

### Thin layer chromatography

*(TLC).* Purified samples of TNP-AMP-PCP (and the products from reactions involving AMP-PNP and ATP-γ-S) were spotted on MilliporeSigma (Burlington, MA) TLC Silica gel 60 F_254_ plates (mobile phase 40:10:25 *n*-butanol: glacial acetic acid: water).

Commercially available TNP-ATP (Tocris Bioscience; Minneapolis, MN) and TNP-ADP (Axxora LLC; Farmingdale, NY) were run for comparison. Retention fraction values were as follows: TNP-ATP, 0; TNP-ADP, 0.006; TNP-AMP-PCP, 0; AMP-PNP product 0.35; ATP-γ-S product (0.5).

### UV-visible spectrophotometry

Aqueous solutions of TNP-AMP-PCP were quantified by absorbance at 259 nm, using the published ε_259 nm_ value for TNP-ATP of 26,400 M-1cm-1 (Hiratsuka and Uchida, 1973). Stock solutions were diluted 1:100 in water and absorbance was measured using a Red Tide UBS650U spectrometer (Ocean Optics; Orlando, FL) and LoggerPro software (Vernier; Beaverton, OR). Trinitrophenylation of the ribose of TNP-AMP- PCP induces the formation of a Meisenheimer complex and two additional absorbance peaks in the visible spectrum (ε_408 nm_ = 26,400 M-1cm-1, ε_470 nm_ = 18,500 M-1cm-1; published values for TNP-ATP). This oxygen atoms connecting the ribose ring to the trinitrophenyl group in this complex can be reversibly protonated with a pK_a_ around 4.5 to yield one of two products (Figure 2C) (Azegami and Iwai, 1975). To study pH effects on the absorbance of the Meisenheimer complex an 8.3 mM stock of TNP-AMP-PCP was diluted 1:200 in 1.0 M tris(hydroxymethyl)aminomethane (Tris), pH 8.0, 1.0 M 4-(2-hydroxyethyl)-1- piperazineethanesulfonic acid (HEPES), pH 7.3, or 0.5 M 2-((morpholino)ethyl)sulfonic acid (MES), pH 5.3. Absorbance spectra were blank subtracted using buffer-only samples.

### Fluorescence emission

The fluorescence emission of a 41.5 µM solution of TNP-AMP-PCP in 1 M Tris, pH 8.0 was measured using a Hitachi F-7000 fluorescence spectrophotometer and FL Solutions software. The samples were excited at 400 nm and emission was measured between 450 nm and 700 nm. Slit widths were 5 nm on the excitation and emission side and the PMT voltage was set to 950 V. Spectra were blank subtracted using a 1.0 M Tris solution.

### High performance liquid chromatography (HPLC)

Sample purity was assessed by reversed-phase HPLC using an Econosphere C_18_ 5µ column (Grace; Columbia, MD). Samples were run on a linear gradient from 0-60% acetonitrile in 0.1 M triethylammonium bicarbonate (TEAB, pH ∼7.5) at a flow rate of 1 mL/min using a Hitachi Elite LaChrome HPLC equipped with an L-2450 diode-array detector. Data were acquired using EZChrom Elite software (Agilent Technologies; Santa Clara, CA).

### Mass spectrometry

TNP-nucleotide samples were diluted to a concentration of 20 μM in 1:1 water:acetonitrile plus 0.1% triethylamine. Samples were analyzed via direct-infusion electrospray ionization using an AB SCIEX 4000 QTRAP mass spectrometer (Framingham, MA). The sample was delivered to the mass spectrometer via a syringe pump operating at a flow rate of 10 µL/min. Electrospray ionization was conducted in negative mode with an inlet spray voltage of -4500 V, a declustering potential of -140 V and a source temperature of 70 °C. Analysis was conducted in enhanced mass spectrometry mode using a dynamic fill time for the ion trap, scanning over a range of 200 to 800 m/z at a rate of 1000 m/z per second. Data were acquired and analyzed using Analyst software (Sciex). Peaks and intensities are reported in Table 1.

**Table 1.** Mass spec peaks.

| m/z | height | m/z | height |
| --- | --- | --- | --- |
| 205.2 | 117790 | 396.96 | 182690 |
| 219.04 | 115380 | 413.04 | 103370 |
| 228 | 983170 | 416.96 | 137020 |
| 228.64 | 146630 | 439.04 | 117790 |
| 228.96 | 204330 | 455.12 | 112980 |
| 229.44 | 127400 | 473.28 | 156250 |
| 229.6 | 146630 | 510.08 | 394230 |
| 230.48 | 117790 | 532 | 192310 |
| 237.12 | 1334100 | 547.92 | 117790 |
| 238.16 | 353370 | 570 | 331730 |
| 239.12 | 257210 | 650 | 514420 |
| 239.68 | 125000 | 651.04 | 149040 |
| 239.92 | 110580 | 705.92 | 100960 |
| 241.2 | 115380 | 712 | 278850 |
| 255.2 | 562500 | 714.96 | 637020 |
| 256.24 | 189900 | 715.92 | 153850 |
| 265.12 | 262020 | 728 | 360580 |
| 274.96 | 132210 | 728.88 | 158650 |
| 283.28 | 1180300 | 737.04 | 235580 |
| 284.24 | 312500 | 743.92 | 180290 |
| 295.12 | 310100 | 752.88 | 312500 |
| 297.2 | 168270 | 758.96 | 144230 |
| 311.2 | 230770 | 774.96 | 300480 |
| 325.2 | 266830 | 790.96 | 173080 |
| 339.2 | 158650 | 796.88 | 158650 |

### ^1^H nuclear magnetic resonance spectroscopy (NMR)

Samples were lyophilized overnight using a FreeZone 2.5 Freeze Dryer (Labconoco, Kansas City, MO) and resuspended in D_2_O at 5-10 mM. Powdered Na_2_CO_3_ was added to increase sample pH, where indicated. Data were acquired using an Avance III 400 MHz NMR (Bruker; Billerica MA) and TopSpin software. Peaks and integrals are reported in Table 2. Peak assignments were based on a published ^1^H-NMR spectrum of TNP-ATP (Stephen et al., 2016).

**Table 2.** ^1^H-NMR peaks and integrals. Boxes are used to group multiplets. Peak f (4.7289 ppm) was not integrated as it was on the shoulder of the H_2_O/HOD peak. We did not integrate the impurity peak around 0 ppm.

| peak | [ppm] | Intensity [abs] | Integral |
| --- | --- | --- | --- |
| h | 8.8499 | 2092430 | 1 |
| h | 8.7465 | 2014814 | 1.004234 |
| a | 8.5334 | 389512 | 1.301907 |
|  | 8.4301 | 3183596 |  |
| b | 8.2622 | 368494 | 1.215862 |
|  | 8.2042 | 3107256 |  |
| c | 6.4821 | 2389398 | 1.129836 |
| d | 5.5431 | 858424 | 1.158591 |
|  | 5.5229 | 1482018 |  |
| e | 5.4762 | 1265976 | 1.168417 |
|  | 5.4534 | 727046 |  |
| f | 4.7289 | 1577966 |  |
| g | 4.2094 | 1644184 | 2.727102 |
|  | 4.1985 | 2497092 |  |
| i | 2.1976 | 1516474 | 2.624964 |
|  | 2.1483 | 2351146 |  |
|  | 2.0978 | 1464712 |  |
|  | 0.0663 | 1039234 |  |

### Molecular biology

*Homo sapiens* Kir6.2 was previously subcloned into pcDNA4/TO using standard techniques. *Homo sapiens* SUR1 was cloned into the mOrange-N1 vector using standard PCR/subcloning techniques, which allows for the expression of protein with a C-terminal mOrange tag. Site directed mutagenesis was performed using the QuikChange kit (Agilent; Santa Clara, CA). All clones were verified by direct sequencing by Azenta-Genewiz (South Plainfield, NJ).

### Cell culture

HEK-293T cells were obtained from and verified by ATCC (Manassas, VA). Frozen stocks were prepared from these cells and stored in liquid nitrogen. No additional verification or mycoplasma testing were performed. New cultures were prepared from frozen stocks after ∼20 passages. Cells were grown in T-75 flasks (Grenier Bio-One; Monroe, NC) in Dulbecco’s Modified Eagle Medium (DMEM; Gibco; Grand Island, NY) supplemented with 10% fetal bovine serum (FBS; Corning; Tewksbury, MA), 100 U/mL penicillin, and 100 µg/mL streptomycin (Gibco). Cultures were maintained at 37 °C in a 5% CO2/95% air atmosphere. Cells were plated in 35 mm dishes (Thermo Scientific; Waltham, MA) on glass coverslips (#1 thickness; Chemglass Life Sciences; Vineland, NJ) a day before transfection.

### Expression of ANAP-tagged proteins in HEK293T cells

ANAP-tagged proteins were obtained as previously described (Chatterjee et al., 2013; Puljung et al., 2019; Puljung, 2021). Briefly, cells were co-transfected with 0.5 µg of Kir6.2 plasmid and 1-1.5 µg of SUR1 plasmid with an amber stop codon (TAG) at the position corresponding to the site at which ANAP was to be inserted. Two additional plasmids were transfected: 1) pANAP, which encodes a tRNA with the appropriate anticodon (CUA) to recognize the amber stop codon and a synthetase enzyme capable of charging the tRNA with ANAP, and 2) peRF1-E55D, encoding a dominant negative eukaryotic exchange factor. TransIT transfection reagent (Mirus; Madison, WI) was used at a ratio of 3 µL/µg of DNA. Just prior to transfection, the medium was replaced with DMEM supplemented with FBS, penicillin/streptomycin, and 20 µM ANAP methyl ester (AsisChem Inc.; Waltham, MA). For electrophysiology experiments, cells were kept at 33 °C after transfection and media was supplemented with 300 µM tolbutamide prepared from a 100 mM stock in DMSO. Experiments were performed 3-5 days post-transfection.

### Unroofing HEK cells for imaging

To gain access to the intracellular face of the plasma membrane, cells were unroofed (Usher et al., 2020a; Heuser, 2000). Briefly, a small fragment of glass coverslip with adherent cells was broken off using jeweler’s forceps. The fragment was dipped in a solution of 0.1% poly-L-lysine for 3x10 s. Cells were unroofed by briefly blotting the coverslip fragment (cell-side down) on a clean piece of filter paper (Cytiva Whatman Grade 1; Wilmington, DE), leaving behind adherent plasma membrane fragments.

### Microscopy/spectroscopy of unroofed HEK cells and cell fragments expressing fluorescently tagged channels

Coverslip fragments were imaged in FluoroDishes (World Precision Instruments; Sarasota FL) filled with bath solution (140 mM KCl, 1 mM ethylenediaminetetraacetic acid (EDTA), 1 mM ethylene glycol-bis(2-aminoethyleether)- N,N,N’,N’-tetraacetic acid (EGTA), 10 mM HEPES, pH7.4). Cells were imaged using a Nikon Ti2E microscope equipped with a 20X air objective (Nikon Plan Fluor ELWD 20X/0.45 NA). Unroofed membranes were imaged using a Nikon TE2000-U (Melville, NY) or Ti2E microscope equipped with a 60X water-immersion objective (Nikon Plan Apo VC, 1.20 NA). ANAP was excited using a ThorLabs LED source (LED4D067; Newton, NJ) with a center wavelength of 385 nm. For imaging ANAP, the filter set contained a 390/18 nm band-pass filter (MF390-18, ThorLabs), an MD416 dichroic mirror (ThorLabs), and a 470/40 nm band-pass filter (MF479-40, ThorLabs). To obtain spectra, the emission filter was replaced with a 400 nm long-pass filter (FEL0400, ThorLabs). To image the mOrange tag, the same light source was used with a broad LED centered at 565 nm. A 530-30 band-pass filter was used for excitation (Chroma; Bellows Falls, VT) with a T550lpxr dichroic mirror (Chroma) and 575/50 band pass emission filter (Chroma).

Images and spectra were collected using an IsoPlane81 camera/spectrograph (Teledyne Princeton Instruments; Princeton, NJ) or IsoPlane160 and coupled PIXIS 256 camera (Teledyne). Acquisition was automated using trigger pulses sent from a Sutter IPA amplifier controlled by SutterPatch software version 2.1.1 (Sutter Instruments; Novato, CA). Exposure times were typically 10 s. Fluorescence emission was passed through a slit and reflected off a grating (150 g/mm groove density, 300 nm blaze wavelength) to produce images for which the x-dimension was replaced with wavelength and the y-dimension retained spatial information (Figure 6A). Spectra were obtained by averaging the intensities at each wavelength over a region of interest. An empty region of the image (no cells or unroofed membranes) was selected and averaged by wavelength to produce a background spectrum, which was subtracted from data from the region of interest. Photobleaching artifacts were corrected as previously described (Puljung et al., 2019; Usher et al., 2020a). Briefly, five spectra were acquired in the absence of nucleotide, the peak intensity of the ANAP spectrum was plotted as a function of exposure time, the intensity decay was fit with a single exponential, and the resulting curve was used to correct subsequent exposures.

### Measuring nucleotide binding

The FluoroDish containing the coverslip with cell fragments was constantly perfused with bath solution using a MiniStar peristaltic pump (World Precision Instruments). Solutions of AMP-PCP, TNP-ATP, and TNP-AMP-PCP were prepared from frozen stocks in bath solution. Stocks and dilutions were stored at -20 °C until use and kept in a covered ice bucket during experiments. Nucleotides were applied directly to the cell fragments using a pressure-driven SmartSquirt Micro-Perfusion System (AutoMate Scientific; Berkeley, CA). The Smart Squirt was controlled using a Sutter IPA amplifier and SutterPatch software. For AMP-PCP/TNP-ATP competition assays, a solution of AMP-PCP alone was applied for 30 s prior to co-application of AMP-PCP and TNP-ATP.

### Electrophysiology

Currents were recorded from inside-out membrane patches excised from HEK293T cells expressing Kir6.2-G334D and SUR1 with ANAP incorporated at position 688 (SUR1-W688ANAP) and a C-terminal mOrange tag. The G334D mutation in Kir6.2 eliminates the inhibitory effect of nucleotides at Kir6.2, allowing us to study activation in isolation (Drain et al., 1998). mOrange and ANAP fluorescence were used to identify cells to patch. Currents were digitized at 10 kHz and long-pass filtered at 1 kHz using a Sutter IPA amplifier and SutterPatch version 2.1.1. Solution exchange was accomplished by gravity. The bath solution contained 107 mM KCl, 2 mM MgCl_2_, 1 mM CaCl_2_, 10 mM EGTA, and 10 mM HEPES. The pH was adjusted to 7.2 with KOH for a final K+ concentration of 140 mM. The sodium salt of adenosine 5’- diphosphate (ADP) was added as indicated. ADP was prepared as a 1 mM stock in bath solution and stored at -20 °C. Pipettes were pulled from borosilicate glass (1B150F-4; World Precision Instruments) to resistances between 1 and 5 MΩ. The pipette solution contained 140 mM KCl, 1.2 mM MgCl_2_, 2.5 mM CaCl_2_, and 10 mM HEPES pH 7.4 with N-methyl-D-glucamine (NMDG). 10 mM Ba^2+^ (10 mM BaCl_2_, 125 mM KCl, 2 mM MgCl_2_, 10 mM HEPES pH 7.2 with KOH) was used to block K_ATP_ current. Raw currents were leak and capacitance corrected for display by subtracting the remaining current in 10 mM Ba^^2+^^.

### Data analysis

Spectra and chromatograms were analyzed and plotted using custom code written in R version 4.5.0, with plotly, hyperSpec, beepr, ggplot2, gridExtra, and xlsx packages (R Core Team; Beleites and Sergo, 2024; Bååth, 2024; Dragulescu and Arendt, 2020; Wickham, 2016; Auguie, 2017; Sievert, 2020). Images were analyzed and displayed using Fiji (Schindelin et al., 2012). Analysis code was deposited on GitHub (https://github.com/puljung/KATP). Data from competition experiments were plotted and fit using custom code written in R version 4.5.0 with the ggplot2 package (R Core Team; Wickham, 2016). Additional plots, hypothesis testing, and curve fits were carried out using GraphPad Prism (Graphpad Software, Inc.; La Jolla, CA) or Microsoft® Excel® for Microsoft 365 MSO (Version 2409 Build 16.0.18025.20160 64 Bit; Redmond WA).

For concentration-response relationships, spectra were acquired at each concentration of nucleotide. ANAP fluorescence was averaged between 460 nm and 470 nm at each concentration and plotted as a function of nucleotide concentration. Fluorescence data were normalized to the lowest concentration. The data were fit with curve of the form

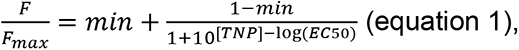

where *F* is fluorescence, *F_max_* is the fluorescence intensity at the lowest nucleotide concentration (where no appreciable quenching was observed), [*TNP*] is the concentration of applied nucleotide, *EC_50_* is the concentration that produced half-maximal quenching, and *min* refers to the residual fluorescence at saturating [*TNP*]. This equation assumes a single binding site with no cooperativity. We obtained better fits when using an equation that incorporated a slope factor. However, in the absence of Mg^2+^, we do not expect nucleotide binding to induce a conformational change in SUR1 that would result in cooperative binding (Puljung et al., 2019).

Thus, we did not believe that the use of an additional free parameter was justified.

In previous work, the curve for TNP-ATP binding to Kir6.2/SUR1-T1397ANAP channels in the absence of Mg^2+^ was fit with the following equation

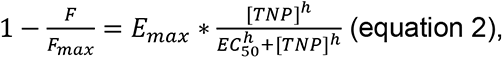

where *E_max_* is the FRET efficiency at saturating TNP-ATP concentrations, *h* is the Hill slope and the other variables have the same meaning as for equation 1 (Puljung et al., 2019). Fits yielded values of *E_max_* = 0.94 ±0.01, *EC_50_* = 4.7 µM ± 0.8 µM, and *h* =0.83.

Data for competition assays between TNP-ATP and non-fluorescent AMP-PCP were fit to the following expression

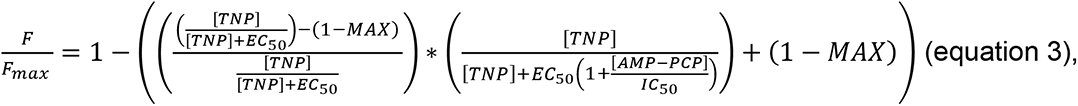

where all variables have their above meanings, [*AMP-PCP*] is the concentration of AMP-PCP, *IC_50_* is the apparent affinity for AMP-PCP, and MAX refers to the maximal asymptote at saturating [AMP-PCP].

Electrophysiology data were analyzed in SutterPatch and Microsoft® Excel® for Microsoft 365 MSO (Version 2409 Build 16.0.18025.20160) 64-bit (Redmond WA).

### Data presentation and statistics

Data are available from the Dryad Digital Repository (DOI: 10.5061/dryad.rv15dv4mw). All error bars represent ±SEM. Wherever possible, all of the individual data points were displayed. *n* values and fit parameters are included in the figure legends. No power analyses were performed prior to conducting experiments to determine the number of experiments. Experimenters were not blinded for any experiments. Means comparisons were conducted using two-tailed Student’s *t*-tests with the threshold of statistical significance at p < 0.01. Protein structures were displayed using ChimeraX developed by the Resource for Biocomputing, Visualization, and Informatics at the University of California, San Francisco with support from National Institutes of Health RO1-GM129325 and the Office of Cyber Infrastructure and Computational Biology, National Institute of Allergy and Infectious Disease (Pettersen et al., 2021). Chemical structures were created and formula masses predicted using ChemDraw (Revvity Signals Software; Waltham, MA).

### Materials and chemicals

All chemicals were obtained from Sigma-Aldrich (St. Louis, MO), unless otherwise noted. pcDNA4/TO was obtained from Invitrogen (Carlsbad, CA). mOrange-N1 was a gift from Michael Davidson (Addgene plasmid # 54499 ; http://n2t.net/addgene:54499 ; RRID:Addgene_54499) (Kremers et al., 2009). pANAP was a gift from Peter Schultz (Addgene plasmid # 48696 ; http://n2t.net/addgene:48696 ; RRID:Addgene_48696) (Chatterjee et al., 2013). peRF1-E55D (*Homo sapiens*) was a kind gift from the Chin Laboratory (MRC Laboratory of Molecular Biology, Cambridge UK) (Schmied et al., 2014).

## RESULTS

### Synthesis and characterization of TNP-AMP-PCP

We adapted an existing protocol for synthesis of trinitrophenyl adenosine and adenosine phosphates to synthesize TNP-AMP-PCP (Figure 2A) (Azegami and Iwai, 1975; Hiratsuka and Uchida, 1973). AMP-PCP was reacted with a four-fold molar excess of TNBS under basic conditions (pH adjusted with LiOH) for four days. The resulting orange product was precipitated with ethanol/sodium acetate and stored at -20 °C. After centrifugation and washing with ethanol, the product was dissolved in water for further analysis.

The UV/Vis spectrum of the product was consistent with expectations based on published spectra of TNP-ATP (Figure 2B) (Hiratsuka and Uchida, 1973). When attached to the ribose of AMP-PCP, the trinitrophenyl group formed a Meisenheimer complex, which was apparent from the appearance of two peaks at 408 nm and 480 nm in addition to the peak at 259 nm, corresponding to the adenine ring (Azegami and Iwai, 1975; Hiratsuka and Uchida, 1973).

Reversible protonation of the Meisenheimer complex results in a loss of these peaks as well as a loss of the orange color (Figure 2C) (Azegami and Iwai, 1975; Hiratsuka and Uchida, 1973). The pK_a_ for this effect in TNP adenosine is around 4.5, so this complex and the color should be present at physiological pH. Consistent with this, the measured UV/Vis spectra in Tris buffer at pH 8.0 and HEPES buffer at pH 7.3 were nearly identical (Figure 2D). Based on published results with TNP-ATP, the ratio of A_259 nm_:A_408 nm_ should be 1:1.06. The A_259 nm_:A_408 nm_ for the product at pH 8.0 was 1:0.99, suggesting contamination from unreacted AMP-PCP (which would only absorb at 259 nm). The A_259 nm_:A_408 nm_ was reduced to 1:0.45 in spectra acquired at pH 5.3, broadly consistent with expectations based on the pK_a_ for TNP-adenosine (Figure 2D).The fluorescence emission spectrum of the product was also measured (Figure 2B). Excitation at 400 nm produced a single emission peak at 551 nm, similar to the value of 561 nm measured for TNP-ATP in water (Hiratsuka, 2003).

Product purity was assessed using reversed-phase HPLC (Figure 2E,F). Samples were run on a C_18_ column on a linear gradient from 0-60% acetonitrile in aqueous 0.1 M triethylammonium bicarbonate buffer (pH 7.5). Using a diode-array detector, a complete UV/Vis spectrum was obtained at every time point. The chromatograms showed one major peak with an absorbance spectrum identical to that expected from TNP-AMP-PCP. The drift in the baseline of the chromatogram at 260 nm, was also observed in blank runs as well and is due to the absorbance of acetonitrile.

Mass spectrometry with electrospray ionization in negative-ion mode showed prominent peaks at m/z ratios of 714.96 Da and 237.12 Da, consistent with the expected values for [TNP-AMP- PCP]- (715 Da) and [TNP-AMP-PCP]3- (237.66 Da) (Figure 3A-C, Table 1). The ratio of intensities of isotope peaks at 716 Da and 717 Da to the intensity of the 715 Da peak was consistent with predictions based on the natural abundances of the constituent elements in TNP-AMP-PCP (Figure 3B). Several peaks at m/z ratios above 715 Da were observed. We assume that these represent various salt adducts of TNP-AMP-PCP and have assigned them where possible. Two of these peaks correspond to the expected mass of TNP-AMP-PCP plus 13 Da and 29 Da, respectively. We could not unequivocally assign these peaks based on their masses but assume them to be Li+ adducts as they were observed in samples of TNP-AMP- PCP (where LiOH was used in the preparation) but not in samples of commercially prepared TNP-ATP, which did not contain Li+.

**Figure 3.**
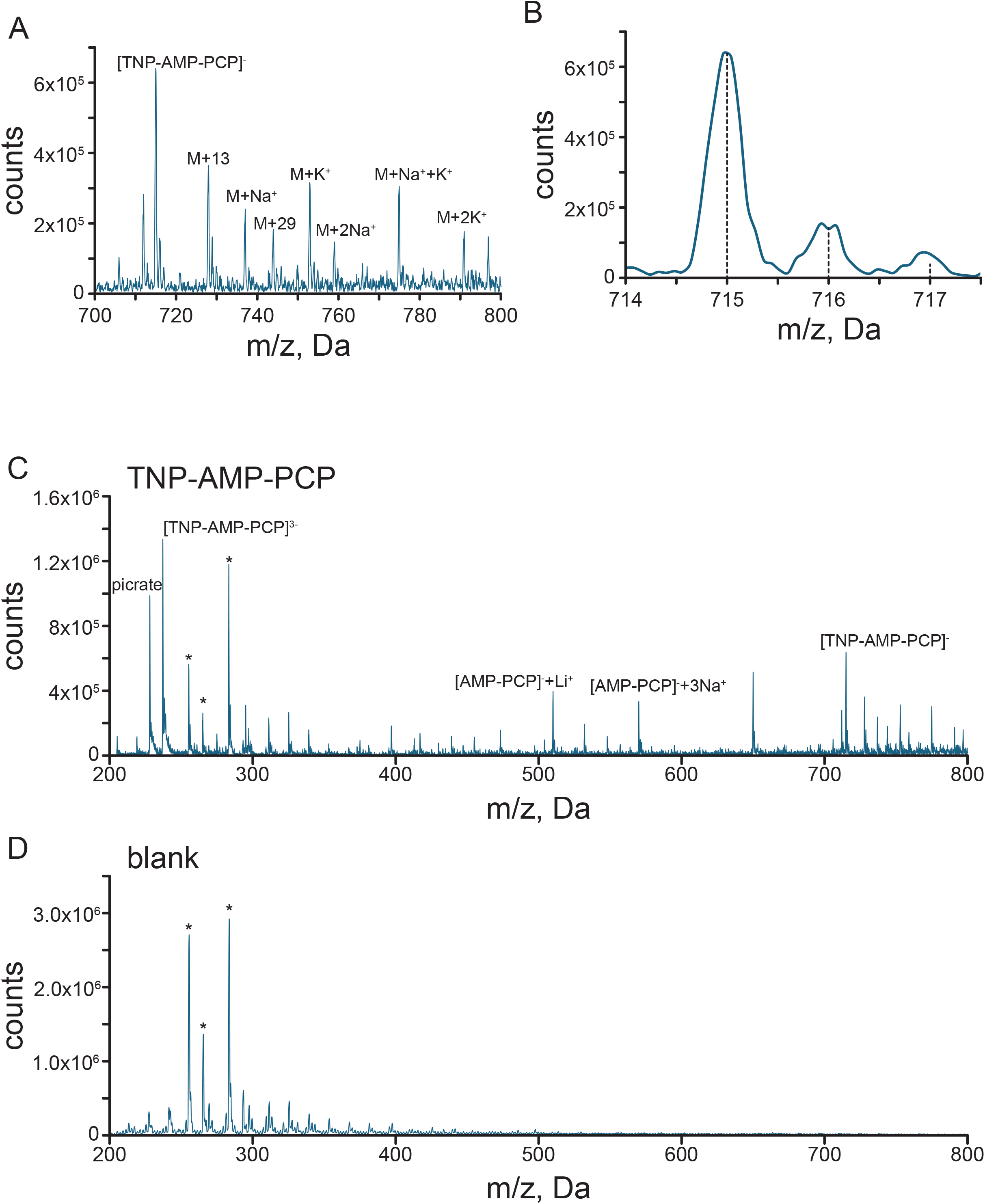
Mass spectrometry of TNP-AMP-PCP. A. Mass spectrum for TNP-AMP-PCP with electrospray ionization in negative-ion mode, showing a peak at the expected mass-to-charged (m/z) ratio (715 Da) for [TNP-AMP-PCP]- as well as several peaks corresponding to likely salt adducts (labeled). B. Zoomed in spectrum showing the three peaks corresponding to isotopic variants of [TNP-AMP-PCP]- and predictions of the expected isotope ratios (dashed lines). C. Mass spectrum showing the complete range examined. Peaks that are also present in the blank samples are marked with “*”. D. Mass spectrum of the blank solution (0.1% triethylamine in 50:50 acetonitrile:water).

Additional peaks between 200-300 Da likely reflected solvent impurities as they showed up in blank measurements (Figure 3C,D; marked by “*”). A substantial peak at 228 Da is likely picrate. Picrate could either be formed by hydrolysis of TNBS, one of the starting materials, or could result from fragmentation of TNP-AMP-PCP, either in solution or in the mass spec. No picrate peak was noted in the HPLC chromatograms, implying it may be a product of fragmentation in the mass spectrometer. Whereas picrate may represent an impurity, based on its structure and spectral properties, it is not expected to interfere with subsequent measurements on K_ATP_.

Finally, ^1^H-NMR was used to verify correct product formation (Figure 4A). The spectrum, obtained in D_2_O, showed peaks consistent with the published structure of TNP-ATP (Stephen et al., 2016). Of note, the peaks at 8.75 and 8.85 ppm correspond to the two protons on the trinitrophenyl group. The triplet centered at 2.15 ppm represents the methylene group between the β and γ phosphates. Some fine splitting of the trinitrophenyl peaks was observed in the original spectrum, which could arise from the protonation of the Meisenheimer complex producing additional structures (Figure 2C,D; Figure 4A,B). Addition of Na_2_CO_3_ to the NMR samples to increase the pH resolved this issue, consistent with this hypothesis (Figure 4B).

**Figure 4.**
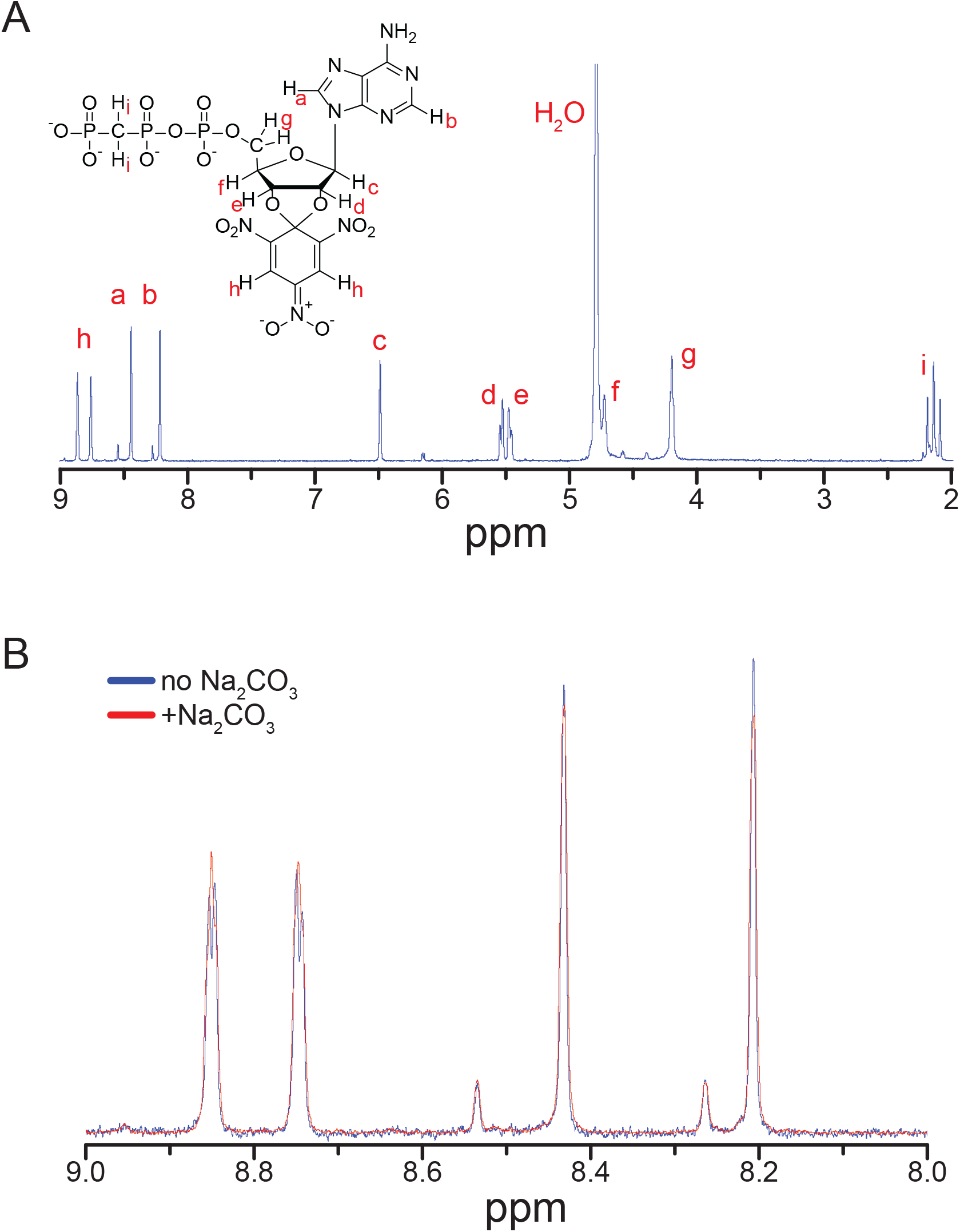
^1^H-NMR spectrum of TNP-AMP-PCP. A. ^1^H-NMR spectrum of TNP-AMP-PCP in D_2_O. Peak assignments are based on the published spectrum of TNP-ATP (Stephen et al., 2016). B. pH dependence of the peaks at 8.7 ppm and 8.8 ppm. The sample was maintained at a basic pH through the addition of solid Na_2_CO_3_.

There was some deviation from expected values in the peak integrals (Table 2). This was likely the result of the large H_2_O/H-OD peak present in the spectra (Figure 4A).

### Measuring nucleotide binding to NBS1 of SUR1

In our previous work, we measured nucleotide binding to Kir6.2 and NBS2 of SUR1 (Usher et al., 2020b; Puljung et al., 2019). However, we made no measurements of nucleotide binding to NBS1. Therefore, we first sought to measure TNP-ATP binding to NBS1. We chose to measure binding in the absence of Mg^2+^ (i.e. in the presence of 1 mM EDTA and no added Mg^2+^). Based on our results at NBS2, we do not expect that Mg^2+^ will be required for binding at either NBS of SUR1. Indeed, it has long been suggested that ATP binding to NBS1 does not require Mg^2+^ (Matsuo et al., 1999). Furthermore, we do not expect TNP-ATP binding to affect the conformation of SUR, as Mg^2+^ is required for NBD dimerization subsequent to nucleotide binding (Puljung et al., 2019).

An amber (TAG) stop codon was introduced into the SUR1 gene at a position corresponding to W688 in the amino acid sequence (SUR1-W688stop, Figure 1C,5A). In the cryo-EM structure of K_ATP_, determined in the presence of MgATP, the tryptophan residue normally occupying this position forms a π-stacking interaction with the adenine ring of ATP (Lee et al., 2017).

**Figure 5.**
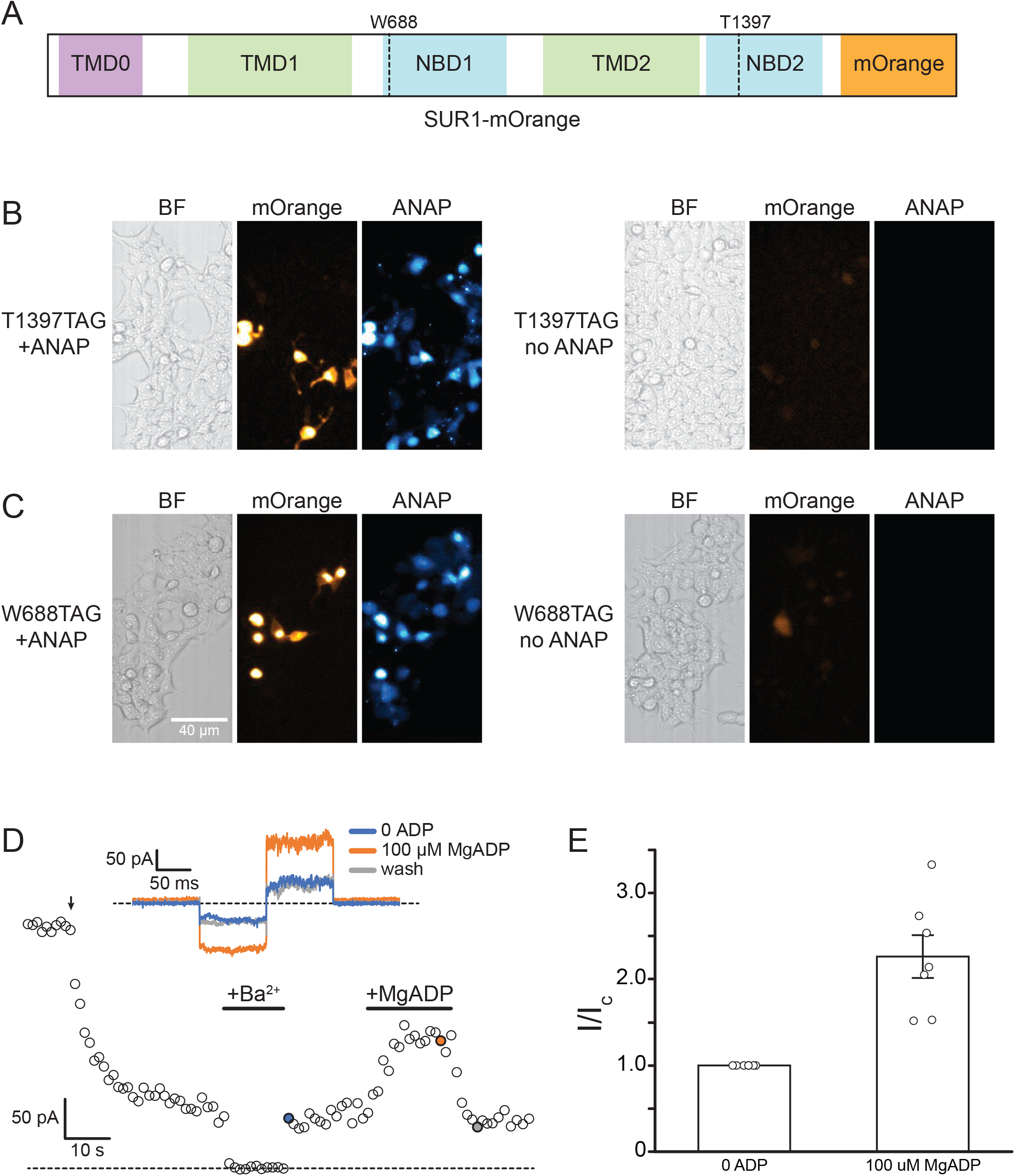
Functional expression of ANAP tagged K_ATP_ channels. A. Schematic diagram of the SUR1-mOrange constructs used showing the location of the two ANAP labeling sites (at positions 688 and 1397). B. *left*. Brightfield and fluorescence images showing HEK293T cells expressing Kir6.2/SUR1-T1397TAG with a C-terminal mOrange tag on SUR1, cultured in the presence of 20 µM ANAP. *right*. Brightfield and fluorescence images showing HEK293T cells expressing Kir6.2/SUR1-T1397TAG C-terminal mOrange tag on SUR1, cultured without ANAP. C. *left,* Brightfield and fluorescence images showing HEK293T cells expressing Kir6.2/SUR1- W688TAG C-terminal mOrange tag on SUR1, cultured in the presence of 20 µM ANAP. *right*. Brightfield and fluorescence images showing HEK293T cells expressing Kir6.2/SUR1-W688TAG C-terminal mOrange tag on SUR1, cultured without ANAP. D. Inward currents from an inside-out patch expressing Kir6.2-G334D/SUR1-W388ANAP with a C-terminal mOrange tag on SUR1. The arrow indicates patch excision. 20 mM Ba^2+^ or 100 µM MgADP were added as indicated. *Inset.* Current traces in response to steps to +20 mV and -20 mV from a holding potential of 0 mV. The colors in the inset correspond to the colored time points in the main figure. E. Activation by 100 µM MgADP. Data were leak corrected by subtracting the current in 20 mM Ba^2+^. ADP increased the current by a factor of 2.3 ± 0.3 (*n* = 7).

**Figure 6.**
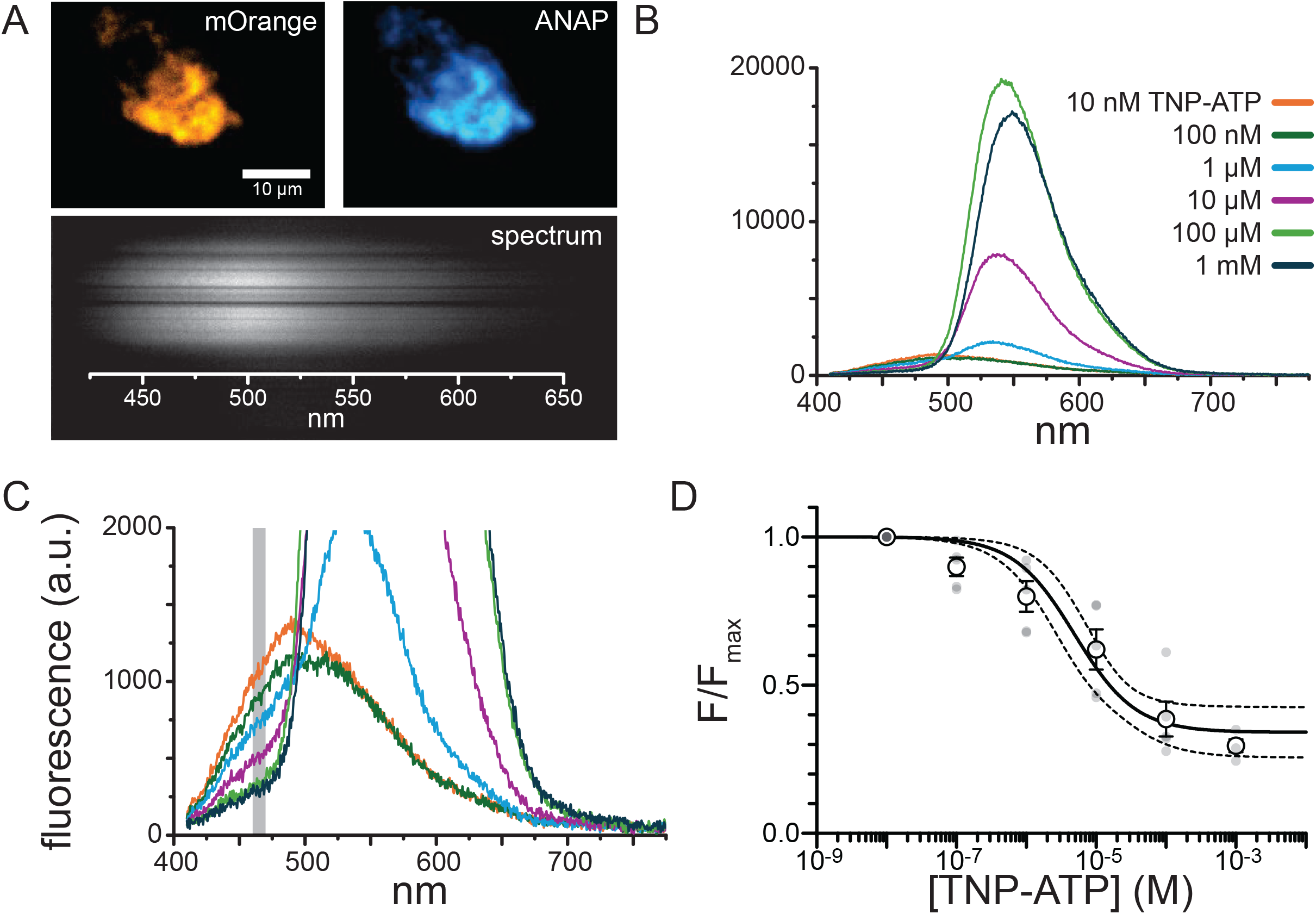
Measuring TNP-ATP binding to NBS1 of SUR1. A. The top panels show mOrange and ANAP fluorescence from an unroofed HEK293T cell expressing Kir6.2/SUR1-W688ANAP channels with a C-terminal mOrange tag. In the bottom panel, the emitted ANAP fluorescence (from the membrane fragment in the epifluorescence image) is reflected off a grating to produce spectra. Note that the x-dimension now represents wavelength, but the y-dimension still represents spatial information (same scale bar as in the top panels). B. Spectra acquired from an unroofed membrane expressing Kir6.2/SUR1-W688 ANAP (with a C-terminal mOrange tag on SUR1) acquired at increasing concentrations of TNP-ATP. The peak around 485 nm corresponds to ANAP and the peak around 551 nm is from TNP-ATP. C. Zoomed in spectra from panel C. The gray bar denotes the wavelength range used to quantify ANAP fluorescence (460-470 nm). D. Concentration-response relationship for ANAP quenching as a function of TNP-ATP concentration. ANAP quenching was quantified by a reduction in fluorescence. Fluorescence intensities were normalized to the ANAP fluorescence at 10 nM TNP-ATP. Data were fit to equation 1. *logEC_50_* = -5.3; *min* = 0.34. Dashed lines indicate 95% confidence intervals for the fit. Individual data points are shown in gray. *n* = 5 for each concentration.

Therefore, ANAP incorporated at this position should be ideally positioned for FRET with bound TNP-ATP. A similar substitution in NBS2 produced robust FRET with TNP-ATP and TNP-ADP (Puljung et al., 2019).

Expressing SUR1-W688stop alone would result in the translation of SUR1 protein truncated after position 687. To express full-length, ANAP-labeled SUR1 proteins, SUR1-W688stop was co-expressed with Kir6.2 and two additional plasmids. The first, pANAP, encodes several copies of a tRNA with the appropriate anticodon (CUA) to recognize the amber stop codon along with a tRNA synthetase capable of charging this tRNA with ANAP (Chatterjee et al., 2013). The second plasmid encodes a dominant negative eukaryotic exchange factor, which helps to increase the yield of full-length, ANAP-tagged channels (Schmied et al., 2014). ANAP was introduced to the cell culture medium before transfection. In past work, 92% full-length, ANAP-labeled channels were obtained under similar conditions (Puljung et al., 2019). To ensure ANAP incorporation, the carboxy terminus of SUR1 was tagged with mOrange. The presence of orange fluorescence along with ANAP fluorescence indicated the production of full-length SUR1-W688ANAP.

Some groups have reported unintended read through of the stop codon, i.e. the production of full-length protein in the absence of ANAP, or reinitiation of translation after a stop codon (Hsieh et al., 2014; Kalstrup and Blunck, 2015; Puljung, 2021). To control for this, SUR1-W688stop and SUR1-T1397stop (ANAP label near NBS2; Figure 1C,5A) were expressed as above, but in the presence and absence of 20 µM ANAP in the cell culture medium (Figure 5B,C). In the presence of ANAP, bright orange fluorescence was noted in cells expressing each construct.

The same cells also showed bright ANAP signals. In the absence of ANAP, strong mOrange signals were not observed for either SUR1 construct. A few cells showed dim, cytoplasmic fluorescence which may correspond to soluble mOrange being transcribed from an internal start site. Nevertheless, the FRET-based measurements reported here all relied on the ANAP signal so full-length channels that lack ANAP would not contribute directly to the measured signal.

Many cells showed diffuse ANAP fluorescence with no concomitant mOrange signal. This background has been widely reported and ascribed to unincorporated ANAP in untransfected cells that took up the dye (Aman et al., 2016; Dai and Zagotta, 2017; Puljung, 2021; Puljung et al., 2019; Usher et al., 2020b; Shandell et al., 2019). As the fluorescence experiments reported here were performed in unroofed plasma membrane fragments, the background from unincorporated, cytosolic ANAP should be negligible. Nevertheless, experiments were performed using cells or cell membranes for which bright ANAP and mOrange signals were observed, thus maximizing the probability that the observations were made from full-length, ANAP labeled channels.

In our previous work, we found that replacing Y1353 in the A-loop of NBD2 of SUR1 with ANAP produced channels that were not activated by MgADP (Puljung et al., 2019). Therefore, we repeated our binding experiments using an alternative ANAP labeling site (T1397ANAP) that produced functional channels that were activated by MgADP. As W688 is in a position in the A loop of NBD1 that is analogous to that of Y1353 in NBD2, we sought to verify that Kir6.2/SUR1- W688ANAP channels were functional. SUR1-W688ANAP was co-expressed with Kir6.2-G334D. The G334D mutation effectively removes the inhibitory effect of nucleotides on Kir6.2 so that activation of channels via SUR1 can be studied in isolation (Drain et al., 1998). Figure 5D shows a representative recording from an inside-out membrane patch excised from an HEK293T cell expressing Kir6.2-G334D/SUR1-W688ANAP. Because Kir6.2-G334D is insensitive to ATP, it produces measurable currents in cell-attached patches. After patch excision (arrow) the current began to run down, presumably due to a loss of phosphatidylinositol-4,5-bisphosphate (Hilgemann and Ball, 1996). The remaining currents were reversibly blocked by the addition of 10 mM Ba^2+^ and reversibly stimulated by 100 µM MgADP. The Ba^2+^-sensitive currents increased by a factor of 2.2 ± 2 (p = 0.003, two-tailed paired t-test). Thus, substitution of W688 with ANAP produced channels that were functional and stimulated by MgADP.

To measure the binding of TNP-ATP to labeled SUR1, one must first expose the cytoplasmic face of the channel. Unroofed membrane fragments were prepared by blotting cells grown on coverslips with filter paper (Heuser, 2000; Usher et al., 2020a). This process removes the cells, leaving behind adherent fragments of plasma membrane with the intracellular face exposed to bulk solution. Unroofed membrane fragments expressing Kir6.2/SUR1-W688ANAP were identified by the presence of fluorescence for both mOrange and ANAP tags (Figure 6A). We assumed the presence of Kir6.2 based on patch-clamp recordings showing functional Kir6.2/SUR1-W688ANAP channels (Figure 5D), and the observation that Kir6.2 and SUR1 must first co-assemble before exiting the endoplasmic reticulum (Zerangue et al., 1999). However, tagging the individual subunits with GFP does allow some degree of independent trafficking of the subunits to the plasma membrane, so it remains possible that some of the signals arise from SUR1 expressed by itself (Makhina and Nichols, 1998).

FRET between ANAP and TNP-ATP was used to measure binding to Kir6.2/SUR1- W688ANAP in unroofed membranes. Fluorescence emission spectra were obtained by passing the emitted light from the unroofed membranes through the slit of a spectrograph mounted on a microscope (Figure 6A,B). This allowed for easy separation of the emission peaks corresponding to ANAP (peak emission around 485 nm) and TNP-ATP (peak emission around 551 nm). There was no apparent mOrange fluorescence in the emission spectra when ANAP was excited at 385 nm, suggesting that any FRET between ANAP and mOrange was negligible. Theoretically, one could measure FRET from the quenching of the donor (ANAP) peak or the sensitization of the acceptor (TNP-ATP peak). However, binding was quantified from the reduction in ANAP fluorescence for two reasons. Firstly, the quenching of the donor peak is directly proportional to the FRET efficiency. Secondly, whereas the ANAP peak is specific to labeled K_ATP_ channels, some fraction of the TNP-ATP fluorescence can be attributed to non-specific TNP-ATP binding to the plasma membrane or incomplete background subtraction (Puljung et al., 2019; Usher et al., 2020a).

The emission spectrum of Kir6.2/SUR1-W688ANAP channels was acquired at concentrations of TNP-ATP ranging from 10 nM to 1 mM (Figure 6B-D). There was some overlap evident between the ANAP peak and the TNP-ATP peaks (Figure 6B). To isolate changes in ANAP emission, quenching was assessed by measuring the average fluorescence over a range from 460 nm to 470 nm (Figure 6C, gray bar). Figure 6D shows the average Kir6.2/SUR1- W688ANAP fluorescence normalized to the fluorescence at 10 nM TNP-ATP, where no quenching was observed. The data were fit with a single-site binding curve (equation 1) with an EC_50_ value of 4.9 μM and a minimum normalized fluorescence of 0.34 at saturating concentrations, indicating 66% quenching (Table 3). We expected a greater degree of quenching between ANAP at position 688 and TNP-ATP bound directly to NBS1, given the proximity and the calculated R_0_ (distance where FRET efficiency is half-maximal) for ANAP/TNP-ATP (about 43 Å) (Puljung et al., 2019). However, the degree of quenching we observed was much greater than the value predicted if TNP-ATP were bound to NBS2 (about 20% FRET) based on the inhibited and apo structures of K_ATP_ (Martin et al., 2017, 2019). The reduced quenching compared to our expectations may reflect TNP-ATP binding in an unfavorable orientation relative to ANAP for FRET to occur or simply insufficient background subtraction in our experiments (due to out of focus fluorescence from nearby structures or inhomogeneities in the field illumination), which could make the baseline fluorescence between 460 nm and 470 nm appear higher.

**Table 3.**
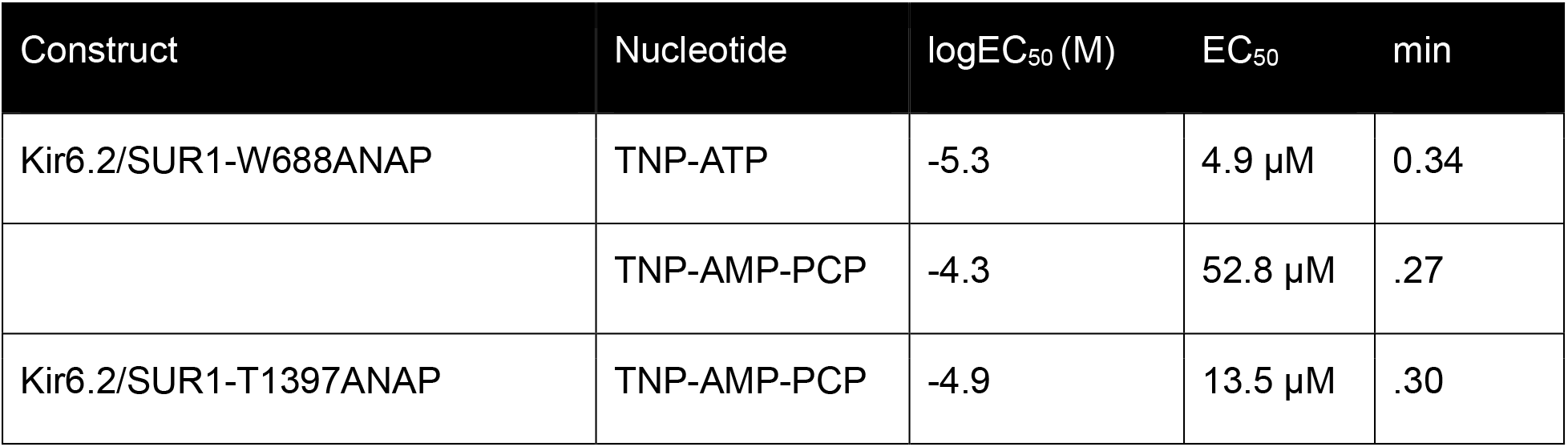
Values from fits to concentration-response curves.

| Construct | Nucleotide | logEC<sub>50</sub> (M) | EC<sub>50</sub> | min |
| --- | --- | --- | --- | --- |
| Kir6.2/SUR1-W688ANAP | TNP-ATP | -5.3 | 4.9 μM | 0.34 |
| | TNP-AMP-PCP | -4.3 | 52.8 μM | .27 |
| Kir6.2/SUR1-T1397ANAP | TNP-AMP-PCP | -4.9 | 13.5 μM | .30 |

### Binding of TNP-AMP-PCP to NBS1 and NBS2

We next tested the ability of our newly synthesized ATP derivative to bind to both NBSs of SUR1 (Figure 7A,B). NBS1 of SUR1 was labeled at position W688. To measure binding to NBS2, the threonine residue at position 1397 was replaced with ANAP (SUR1-T1397ANAP, Figure 1C,5A). In previous experiments, we measured TNP-ATP and TNP-ADP binding at this site in the presence and absence of Mg^2+^ (Puljung et al., 2019). Furthermore, Kir6.2-G334D/SUR1-T1397ANAP channels (in which the inhibitory site on Kir6.2 was mutated) were functional and activated by MgADP (Puljung et al., 2019).

**Figure 7.**
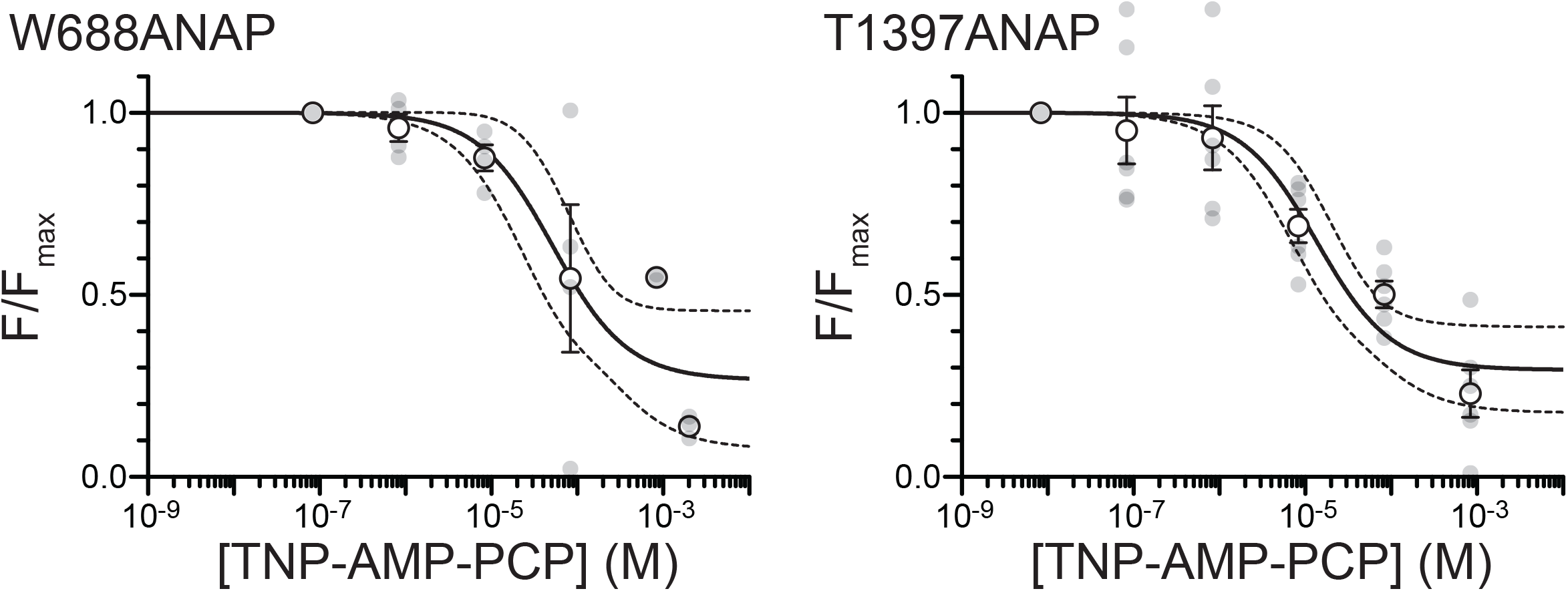
Binding of TNP-AMP-PCP to NBS1 and NBS2 of SUR1. Curves showing concentration-dependent quenching of Kir6.2/SUR1-W688ANAP (NBS1 labeled) and Kir6.2/SUR1-T1397ANAP (NBS2 labeled) by TNP-AMP-PCP. Both SUR1 constructs had C- terminal mOrange tags. Data were fit with equation 1. Dashed lines indicate 95% confidence intervals for the fit. Individual data points are shown in gray. Kir6.2/SUR1-W688ANAP: *logEC_50_* = -4.3; *min* = 0.26. A total of 7 membranes were used in these experiments. The *n* values at each concentration are as follows: 83 nM, *n* = 7; 830 nM, *n* = 4; 8.3 µM, *n =* 4; 83 µM, *n* = 4, 830 µM, *n* = 2; 2 mM, *n* = 3. Kir6.2/SUR1-T1397ANAP: *logEC_50_* = -4.9; *min* = 0.29. A total of 6 membranes were used in these experiments. *n* = 6 for each concentration.

Concentration-response relationships for each NBS were fit to single-site binding curves. For NBS1 (Kir6.2/SUR1-W688ANAP), half-maximal quenching occurred at a concentration of 52.8 μM TNP-AMP-PCP. ANAP fluorescence was quenched by 73% at saturating TNP-AMP-PCP, again consistent with direct binding to NBS1 (Figure 7A, Table 3). For NBS2 (Kir6.2/SUR1- T1397ANAP), half-maximal quenching occurred at 13.5 μM TNP-AMP-PCP. ANAP fluorescence was quenched by 70% at saturating concentrations (Figure 7B, Table 3). By comparison, TNP-ATP quenching of Kir6.2/SUR1-T1397ANAP had an EC_50_ value of 4.7 μM, with 94% quenching at saturating concentrations (Puljung et al., 2019). Based on these data, we conclude that TNP- AMP-PCP was able to bind both NBSs of SUR1.

However, the ability of TNP-AMP-PCP to bind SUR1 does not guarantee that non-fluorescent AMP-PCP is also able to bind. For example, it is possible that the trinitrophenyl moiety was able to stabilize binding of the TNP-AMP-PCP to SUR1, and that AMP-PCP would not appreciably bind without it. As a complementary means of assessing AMP-PCP binding to SUR1, we tested its ability to compete with TNP-ATP for binding at NBS2 of SUR1 (Figure 8). In the absence of AMP-PCP, 5 μM TNP-ATP quenched the fluorescence of Kir6.2/SUR1-T1397ANAP by 44% ± 9%, close to the expected value of 48% based on previous experimental data (Figure 8A,C) (Puljung et al., 2019). However, when the unroofed membrane fragments were perfused with 100 μM AMP-PCP before co-application of 100 μM AMP-PCP and 5 μM TNP-ATP, the quenching was reduced to 22% ± 1% (Figure 8B,C). The difference in quenching was statistically significant (p < 0.01) and implies direct binding of AMP-PCP to NBS2 of SUR1. We further examined the quenching by 5 µM TNP-ATP at a range of AMP-PCP concentrations (Figure 8D). The data were initially fit to a model that assumed a single nucleotide binding site. This model failed to fit the data, as it assumes that the normalized fluorescence would be 1 at saturating concentrations of AMP-PCP, which was inconsistent with the data. Instead, the data were fit with a model assuming competition for a single site, but with an offset at saturating AMP-PCP (equation 3). This model had three free parameters: offset, apparent TNP-ATP affinity and apparent AMP-PCP affinity. The fits returned a value of 5.31 µM for TNP-ATP apparent affinity, consistent with the published value of 4.7 µM (Puljung et al., 2019). Regardless, the overall quality of the fit was poor. Whereas there was an overall concentration-dependence to the competition between the two nucleotides, saturating concentrations of AMP-PCP did not produce a complete recovery of the Kir6.2/SUR1-T1397ANAP fluorescence (as would be expected for competition at a single binding site). The residual quenching at high AMP-PCP concentrations may reflect TNP-ATP binding to another site on KATP (like the one on Kir6.2). As this relationship did not follow predictions for competition at a single binding site and fits to equation 3 were poor, a quantitative interpretation of these results is not possible.

**Figure 8.**
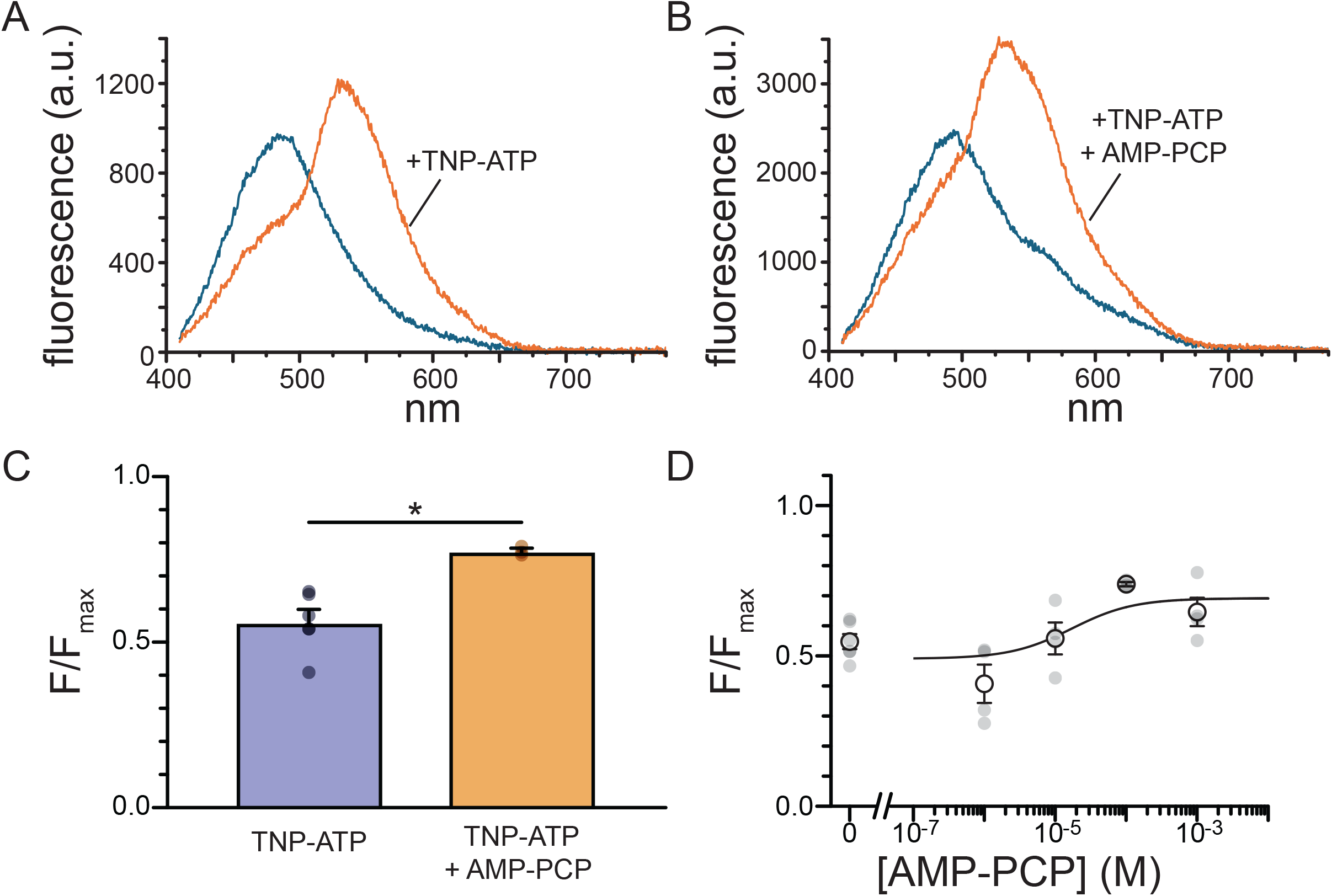
AMP-PCP binds to NBS2 of SUR1. A. Fluorescence of Kir6.2/SUR1-T1397ANAP in the presence and absence of 5 µM TNP-ATP. B. Fluorescence of Kir6.2/SUR1-T1397ANAP in the presence and absence of 5 µM TNP-ATP plus 100 µM AMP-PCP. C. Bar chart summarizing the reduction in Kir6.2/SUR1-T1397ANAP fluorescence in 5 µM TNP-ATP in the absence and presence of 100 µM AMP-PCP. *n* = 6 for 5 µM TNP-ATP; *n* = 3 for 5 µM TNP-ATP plus 100 µM TNP-ATP. Individual data points are superimposed on the graph. Data sets were compared using a two-tailed Student’s *t*-test. * indicates p < 0.01. D. Kir6.2/SUR1-T1397ANAP fluorescence in 5 µM TNP-ATP as a function of AMP-PCP concentration. Data were fit with equation 3. *EC_50_* = 5.31 µM, *IC_50_*= 8.82 µM, *MAX* = 0.73. *n* = 6 for 0 AMP-PCP; *n* = 4 for 1 µM; *n* = 4 for 10 µM; *n* = 4 for 1 µM; *n* = 3 for 100 µM; *n* = 4 for 1 µM; *n* = 4 for 1 mM. Note the broken axis used to show the 0 AMP-PCP point.

## DISCUSSION

We have successfully synthesized and purified TNP-AMP-PCP, a fluorescent, non-hydrolyzable ATP analog. Using our established, FRET-based approach, we measured binding of this analog to both NBSs of SUR1. We also measured the binding of TNP-ATP to NBS1, which had not been examined previously. Finally, we demonstrated that AMP-PCP can bind to NBS2 of SUR1 using a competition assay.

ATP binding to NBS1 was previously observed in cryo-EM structures and in equilibrium binding experiments performed in the presence and absence of Mg^2+^ (Matsuo et al., 1999; Lee et al., 2017). Whereas the presence of TNP-ATP/TNP-ADP at NBS1 could be inferred from our previous studies on NBS2, we did not previously make any direct measurements of nucleotide binding to NBS1. In this study, we used our assay to measure direct TNP-ATP binding to NBS1 in the absence of Mg^2+^ with apparent affinity of 4.9 µM (Figure 6). This is similar to the apparent affinity previously reported for TNP-ATP binding to NBS2 (Kir6.2/T1397ANAP, 4.7 µM) (Puljung et al., 2019). It has been hypothesized that, for asymmetric ABC transporters like CFTR and TM287/288, the degenerate NBS (NBS1 here) binds nucleotides with a very high affinity and that bound nucleotides may act as a structural support, anchoring the NBDs together (Stockner et al., 2020; Hohl et al., 2012). Based on these results, it appears that this may not be true for K_ATP_ channels. This is consistent with existing cryo-EM structures of K_ATP_ that show the NBDs either tightly bundled in the presence of Mg^2+^ and nucleotides or completely separated (Driggers and Shyng, 2023). However, it should be noted that the experiments presented here were performed using a nucleotide derivative and the affinity of this site for unlabeled ATP was not directly tested. Indeed, equilibrium binding studies suggest that ATP can reside at NBS1 of SUR1 for long periods of time (Matsuo et al., 1999).

We report here the first synthesis of TNP-AMP-PCP, a fluorescent non-hydrolyzable ATP analog. Based on UV/Vis and HPLC analyses, a relatively pure sample of TNP-AMP-PCP was obtained (Figure 2E,F). Mass spectrometry showed the presence of some impurities that were not obvious from the HPLC traces (Figure 3). The peak at 228 Da was likely picrate, a potential fragmentation product of TNP-AMP-PCP. However, based on the ratio of the peaks in the UV/Vis spectra, no significant loss of the trinitrophenyl group from the TNP-AMP-PCP samples was observed prior to mass spectrometry. Another possible source of picrate is the hydrolysis of TNBS, which was present in a four-fold excess in the synthesis, but no picrate peak was evident in the HPLC traces. Regardless, if picrate were a substantial contaminant in the purified TNP- AMP-PCP samples, one would not expect it to interfere with subsequent measurements, as 1) it is unlikely to bind to SUR1 and 2) if it were to bind, it would not FRET with ANAP. Likewise, whereas a substantial AMP-PCP contamination is unlikely, such an impurity would compete with TNP-AMP-PCP for binding to SUR1. This would affect the estimated EC50 values, it would not undermine the ultimate conclusion that TNP-AMP-PCP can bind to both NBSs.

Importantly, use of TNP-AMP-PCP allowed for the real-time measurement of binding to K_ATP_ channels in the plasma membrane. Based on the results presented here, non-hydrolyzable ATP analogs can bind both NBS1 and NBS2 of SUR1, albeit at relatively low apparent affinity. The experiments produced estimates for TNP-AMP-PCP apparent affinity of around 50 µM for NBS1 and 13 µM for NBS2. Absent any quantitative measurement of AMP-PCP affinity, one can, nevertheless, conclude that AMP-PCP competes for TNP-ATP binding at NBS2. Thus, the inability of non-hydrolyzable ATP analogs to activate K_ATP_ is not due to a failure to bind.

Ortiz *et al*. (Ortiz et al., 2013) previously examined the ability of non-hydrolyzable nucleotides to bind SUR1. In their experiments, binding was assessed by the ability of nucleotides to displace tritiated glibenclamide from isolated SUR1 subunits expressed in *Pichia* membranes in equilibrium binding assays. ATP-γ-S was able to displace glibenclamide binding, implying that it not only bound to SUR1, but induced it to adopt an “active” conformation. In contrast to this, AMP-PNP and AMP-PCP did not displace glibenclamide but did reduce the ability of ATP to displace glibenclamide, implying that they compete with ATP for binding. Indeed, the authors further demonstrated that AMP-PCP could displace 8-azido-[α-32P]ATP from NBS1. Bernardi *et al*. also showed that AMP-PNP and ATP-γ-S can both compete for α-[32P]Ox-ATP binding to SUR subunits purified from pig brain (Bernardi et al., 1988).

The most direct readout of nucleotide effects on K_ATP_ activation comes from inside-out patch-clamp experiments in which the experimenter can easily change the contents of the “intracellular” bathing solution. For the majority of such experiments, authors report only inhibition by AMP-PCP, AMP-PNP, and ATP-γ-S in the presence of Mg^2+^, which is mediated by binding to Kir6.2 (Ashcroft and Kakei, 1989; Treherne and Ashford, 1992; Jiang and Haddad, 1997; Schwanstecher and Panten, 1994; Schwanstecher et al., 1994, 1992). However, it can be difficult to isolate stimulatory nucleotide effects in wild-type channels with intact inhibition at Kir6.2. For native Kir6.2/SUR1 channels in β-cells and heterologously expressed in *Xenopus* oocytes, the stimulatory effects of SUR1 in the presence of ATP manifest as a shift of the inhibition curve to higher concentrations of ATP in the presence of Mg^2+^ (Ashcroft and Kakei, 1989; Proks et al., 2010). Hehl and Neumcke reported a similar rightward shift in the concentration response curve for AMP-PNP in the presence of Mg^2+^ for K_ATP_ channels in inside-out patches from mouse skeletal muscle, consistent with mixed activation and inhibition by AMP-PNP (Hehl and Neumcke, 1994). However, no such Mg^2+^ effect was observed in inside-out patches from pancreatic β-cells (Schwanstecher et al., 1992). This may reflect differences in K_ATP_ subtypes between the two tissues (Kir6.2/SUR1 in β-cells and Kir6.2/SUR2A in muscle) (Ashcroft, 2023). In each case, concentrations ranged from tens of µM to low mM, the range over which we observed TNP-AMP-PCP binding to both NBSs of SUR1 (Figure 7).

The most direct way to measure nucleotide activation is to measure the response of channels bearing a point mutation that disrupts the inhibitory NBS on Kir6.2 (Kir6.2-G334D) in inside-out patches (Drain et al., 1998; Li et al., 2002; Proks et al., 2010, 2014). Proks *et al*. measured robust activation by MgATP-γ-S of Kir6.2-G334D/SUR1 in inside-out patches from *Xenopus* oocytes channels, but observed no such activation by MgAMP-PNP (Proks et al., 2010). To our knowledge, no such experiments have been performed with AMP-PCP.

The experiments presented here have the potential to build on previous results in significant ways. First of all, the experimental approach allowed for the measurement of nucleotide binding in real time in intact, functional K_ATP_ channel complexes in the plasma membrane of mammalian cells. As such, this method is fully compatible with patch clamp, allowing binding and channel activity to be measured simultaneously (Usher et al., 2020b). Finally, the short distance dependence of FRET allowed the measurement of binding in a site-specific fashion so that the effects of the NBSs could be examined independently.

The conclusion, that non-hydrolyzable ATP analogs bind both NBS1 and NBS2 of SUR1 represents only the first step in understanding the requirement, or lack thereof, for SUR1 to hydrolyze ATP to ADP in order to activate K_ATP_. At least two possibilities remain to explain the inability of non-hydrolyzable ATP analogs to support channel activation. The first possibility is that, while able to bind, non-hydrolyzable ATPs do not stabilize a conformation in SUR1 that promotes activation (NBD dimerization). The second possibility is that hydrolysis of ATP to ADP is a requirement for K_ATP_ activation via SUR1. The ability of AMP-PCP to allow TM287/288 to adopt an outward-facing conformation raises the intriguing possibility that non-hydrolyzable nucleotides may support NBD dimerization in SUR1, but not increase the open probability of the channel’s pore (Hohl et al., 2012). In previous work, we noted that nucleotide dissociation from NBS2 was greatly slowed by the presence of Mg^2+^ (Puljung et al., 2019). This was interpreted to mean that NBD dimerization in the presence of Mg^2+^ impeded nucleotide dissociation. This metric can be used as an indicator of the ability of TNP-AMP-PCP to promote an activating conformational change in SUR1. Again, a key advantage of this technique is that one can also perform such measurements using patch-clamp fluorometry, allowing experimenters to monitor changes in channel and conformational changes in SUR1 simultaneously. When combined with more traditional structure-function approaches such experiments may help to delineate the role of ATP hydrolysis in the gating cycle of K_ATP_. This technique may be further adapted to study the enzymatic activity of other ABC transporters, P-type ATPases, and kinases.

## ACKNOWLEDGMENTS

We would like to thank Dr. Ysobel Baker, Dr. Cheyenne Brindle and Dr. Timothy Curran for helpful discussions and Jazmin Johnson and Christina Alcaro for additional technical support. Research reported in this publication was supported by the National Institute of General Medical Sciences of the National Institutes of Health under award number R15GM155848.

## AUTHOR CONTRIBUTIONS

PR, SW, JA, and MP performed the experiments. SW, JA, and MP performed the data analysis. JA and MP wrote the manuscript.

## Notes

### Competing Interest Statement

The authors have declared no competing interest.

### Summary of Updates

Updated with additional text and revised Figure 1.

## REFERENCES

1. Aguilar-Bryan, L., C. Nichols, S. Wechsler, J. Clement, A. Boyd, G. Gonzalez, H. Herrera-Sosa, K. Nguy, J. Bryan, and D. Nelson. 1995. Cloning of the beta cell high-affinity sulfonylurea receptor: a regulator of insulin secretion. Science. 268:423–426. doi:10.1126/science.7716547.

2. Aleksandrov, L., A.A. Aleksandrov, X.-B. Chang, and J.R. Riordan. 2002. The First Nucleotide Binding Domain of Cystic Fibrosis Transmembrane Conductance Regulator Is a Site of Stable Nucleotide Interaction, whereas the Second Is a Site of Rapid Turnover. J Biol Chem. 277:15419–15425. doi:10.1074/jbc.M111713200.

3. Aman, T.K., S.E. Gordon, and W.N. Zagotta. 2016. Regulation of CNGA1 Channel Gating by Interactions with the Membrane. J. Biol. Chem. 291:9939–9947. doi:10.1074/jbc.M116.723932.

4. Ashcroft, F.M. 2023. KATP Channels and the Metabolic Regulation of Insulin Secretion in Health and Disease: The 2022 Banting Medal for Scientific Achievement Award Lecture. Diabetes. 72:693–702. doi:10.2337/dbi22-0030.

5. Ashcroft, F.M., and M. Kakei. 1989. ATP-sensitive K+ channels in rat pancreatic beta-cells: modulation by ATP and Mg^2+^ ions. J Physiol. 416:349–367. doi:10.1113/jphysiol.1989.sp017765.

6. Ashcroft, F.M., M.C. Puljung, and N. Vedovato. 2017. Neonatal Diabetes and the K ATP Channel: From Mutation to Therapy. Trends in Endocrinology & Metabolism. 28:377–387. doi:10.1016/j.tem.2017.02.003.

7. Auguie, B. 2017. _gridExtra: Miscellaneous Functions for “Grid” Graphics_. R package version 2.3.

8. Azegami, M., and K. Iwai. 1975. Trinitrophenylation of Nucleic Acids and Their Constituents. The Journal of Biochemistry. 78:409–420. doi:10.1093/oxfordjournals.jbchem.a130921.

9. Bååth, R. 2024. _beepr: Easily Play Notification Sounds on any Platform_. R package version 2.0,.

10. Beleites, C., and V. Sergo. 2024. hyperSpec: a package to handle hyperspectral data sets in R’, R package version 0.100.2.

11. Bernardi, H., M. Fosset, and M. Lazdunski. 1988. Characterization, purification, and affinity labeling of the brain [3H]glibenclamide-binding protein, a putative neuronal ATP- regulated K+ channel. Proceedings of the National Academy of Sciences. 85:9816– 9820. doi:10.1073/pnas.85.24.9816.

12. Chatterjee, A., J. Guo, H.S. Lee, and P.G. Schultz. 2013. A Genetically Encoded Fluorescent Probe in Mammalian Cells. J. Am. Chem. Soc. 135:12540–12543. doi:10.1021/ja4059553.

13. Choi, K.-H., M. Tantama, and S. Licht. 2008. Testing for Violations of Microscopic Reversibility in ATP-Sensitive Potassium Channel Gating. J. Phys. Chem. B. 112:10314–10321. doi:10.1021/jp712088v.

14. Csanady, L., P. Vergani, and D.C. Gadsby. 2010. Strict coupling between CFTR’s catalytic cycle and gating of its Cl- ion pore revealed by distributions of open channel burst durations. Proceedings of the National Academy of Sciences. 107:1241–1246. doi:10.1073/pnas.0911061107.

15. Dai, G., and W.N. Zagotta. 2017. Molecular mechanism of voltage-dependent potentiation of KCNH potassium channels. eLife. 6:e26355. doi:10.7554/eLife.26355.

16. Dragulescu, A., and C. Arendt. 2020. _xlsx: Read, Write, Format Excel 2007 and Excel 97/2000/XP/2003 Files_. R package version 0.6.5.

17. Drain, P., L. Li, and J. Wang. 1998. KATP channel inhibition by ATP requires distinct functional domains of the cytoplasmic C terminus of the pore-forming subunit. Proc Natl Acad Sci U S A. 95:13953–13958. doi:10.1073/pnas.95.23.13953.

18. Driggers, C.M., and S.-L. Shyng. 2023. Mechanistic insights on KATP channel regulation from cryo-EM structures. J Gen Physiol. 155:e202113046. doi:10.1085/jgp.202113046.

19. Dunne, M.J. 1989. Protein phosphorylation is required for diazoxide to open ATP-sensitive potassium channels in insulin (RINm5F) secreting cells. FEBS Letters. 250:262–266. doi:10.1016/0014-5793(89)80734-3.

20. Findlay, I. 1987. ATP-sensitive K+ channels in rat ventricular myocytes are blocked and inactivated by internal divalent cations. Pflugers Arch. 410:313–320. doi:10.1007/BF00580282.

21. Gribble, F.M., S.J. Tucker, T. Haug, and F.M. Ashcroft. 1998. MgATP activates the cell KATP channel by interaction with its SUR1 subunit. Proceedings of the National Academy of Sciences. 95:7185–7190. doi:10.1073/pnas.95.12.7185.

22. Hehl, S., and B. Neumcke. 1994. KATP channels of mouse skeletal muscle: mechanism of channel blockage by AMP-PNP. Eur Biophys J. 23:231–237. doi:10.1007/BF00213573.

23. Heuser, J. 2000. The Production of ‘Cell Cortices’ for Light and Electron Microscopy. Traffic. 1:545–552. doi:10.1034/j.1600-0854.2000.010704.x.

24. Hilgemann, D.W., and R. Ball. 1996. Regulation of Cardiac Na+ ,Ca^2+^ Exchange and K_ATP_ Potassium Channels by PIP_2_. Science. 273:956–959. doi:10.1126/science.273.5277.956.

25. Hiratsuka, T. 2003. Fluorescent and colored trinitrophenylated analogs of ATP and GTP. European Journal of Biochemistry. 270:3479–3485. doi:10.1046/j.1432-1033.2003.03748.x.

26. Hiratsuka, T., and K. Uchida. 1973. PREPARATION AND PROPERTIES OF 2’(or Y)-O-(2,4,6- TRINITROPHENYL) ADENOSINE 5’-TRIPHOSPHATE, AN ANALOG OF ADENOSINE TRIPHOSPHATE. Biochimica et Biophysical Acta (BBA) - General Subjects. 320:635–647.

27. Hohl, M., C. Briand, M.G. Grütter, and M.A. Seeger. 2012. Crystal structure of a heterodimeric ABC transporter in its inward-facing conformation. Nat Struct Mol Biol. 19:395–402. doi:10.1038/nsmb.2267.

28. Hsieh, T., N.B. Nillegoda, J. Tyedmers, B. Bukau, A. Mogk, and G. Kramer. 2014. Monitoring Protein Misfolding by Site-Specific Labeling of Proteins In Vivo. PLoS ONE. 9:e99395. doi:10.1371/journal.pone.0099395.

29. Inagaki, N., T. Gonoi, J.P. Clement, N. Namba, J. Inazawa, G. Gonzalez, L. Aguilar-Bryan, S. Seino, and J. Bryan. 1995. Reconstitution of *I*_KATP_ : An Inward Rectifier Subunit Plus the Sulfonylurea Receptor. Science. 270:1166–1170. doi:10.1126/science.270.5239.1166.

30. Inagaki, N., T. Gonoi, and S. Seino. 1997. Subunit stoichiometry of the pancreatic β-cell ATP- sensitive K + channel. FEBS Letters. 409:232–236. doi:10.1016/S0014-5793(97)00488-2.

31. Jiang, C., and G.G. Haddad. 1997. Modulation of K+ channels by intracellular ATP in human neocortical neurons. J Neurophysiol. 77:93–102. doi:10.1152/jn.1997.77.1.93.

32. Kalstrup, T., and R. Blunck. 2015. Reinitiation at non-canonical start codons leads to leak expression when incorporating unnatural amino acids. Sci Rep. 5:11866. doi:10.1038/srep11866.

33. Kozlowski, R.Z., and M.L. Ashford. 1992. Nucleotide-dependent activation of KATP channels by diazoxide in CRI-G1 insulin-secreting cells. Br J Pharmacol. 107:34–43. doi:10.1111/j.1476-5381.1992.tb14460.x.

34. Kremers, G.-J., K.L. Hazelwood, C.S. Murphy, M.W. Davidson, and D.W. Piston. 2009. Photoconversion in orange and red fluorescent proteins. Nat Methods. 6:355–358. doi:10.1038/nmeth.1319.

35. Lee, K.P.K., J. Chen, and R. MacKinnon. 2017. Molecular structure of human KATP in complex with ATP and ADP. eLife. 6:e32481. doi:10.7554/eLife.32481.

36. Li, L., X. Geng, and P. Drain. 2002. Open State Destabilization by Atp Occupancy Is Mechanism Speeding Burst Exit Underlying KATP Channel Inhibition by Atp. The Journal of General Physiology. 119:105–116. doi:10.1085/jgp.119.1.105.

37. Makhina, E.N., and C.G. Nichols. 1998. Independent trafficking of KATP channel subunits to the plasma membrane. J Biol Chem. 273:3369–3374. doi:10.1074/jbc.273.6.3369.

38. Martin, G.M., B. Kandasamy, F. DiMaio, C. Yoshioka, and S.-L. Shyng. 2017. Anti-diabetic drug binding site in a mammalian KATP channel revealed by Cryo-EM. eLife. 6:e31054. doi:10.7554/eLife.31054.

39. Martin, G.M., M.W. Sung, Z. Yang, L.M. Innes, B. Kandasamy, L.L. David, C. Yoshioka, and S.- L. Shyng. 2019. Mechanism of pharmacochaperoning in a mammalian KATP channel revealed by cryo-EM. eLife. 8:e46417. doi:10.7554/eLife.46417.

40. Matsuo, M., N. Kioka, T. Amachi, and K. Ueda. 1999. ATP Binding Properties of the Nucleotide- binding Folds of SUR1. Journal of Biological Chemistry. 274:37479–37482. doi:10.1074/jbc.274.52.37479.

41. Matsuo, M., K. Tanabe, N. Kioka, T. Amachi, and K. Ueda. 2000. Different Binding Properties and Affinities for ATP and ADP among Sulfonylurea Receptor Subtypes, SUR1, SUR2A, and SUR2B. Journal of Biological Chemistry. 275:28757–28763. doi:10.1074/jbc.M004818200.

42. Ortiz, D., L. Gossack, U. Quast, and J. Bryan. 2013. Reinterpreting the Action of ATP Analogs on KATP Channels. Journal of Biological Chemistry. 288:18894–18902. doi:10.1074/jbc.M113.476887.

43. Ortiz, D., P. Voyvodic, L. Gossack, U. Quast, and J. Bryan. 2012. Two Neonatal Diabetes Mutations on Transmembrane Helix 15 of SUR1 Increase Affinity for ATP and ADP at Nucleotide Binding Domain 2. Journal of Biological Chemistry. 287:17985–17995. doi:10.1074/jbc.M112.349019.

44. Pettersen, E.F., T.D. Goddard, C.C. Huang, E.C. Meng, G.S. Couch, T.I. Croll, J.H. Morris, and T.E. Ferrin. 2021. UCSF ChimeraX: Structure visualization for researchers, educators, and developers. Protein Sci. 30:70–82. doi:10.1002/pro.3943.

45. Proks, P., H. de Wet, and F.M. Ashcroft. 2010. Activation of the KATP channel by Mg-nucleotide interaction with SUR1. Journal of General Physiology. 136:389–405. doi:10.1085/jgp.201010475.

46. Proks, P., H. de Wet, and F.M. Ashcroft. 2014. Sulfonylureas suppress the stimulatory action of Mg-nucleotides on Kir6.2/SUR1 but not Kir6.2/SUR2A KATP channels: A mechanistic study. Journal of General Physiology. 144:469–486. doi:10.1085/jgp.201411222.

47. Puljung, M. 2023. KATP channels and the regulation of insulin secretion. *In* TEXTBOOK OF ION CHANNELS: three volume set. J. Zheng and M.C. Trudeau, editors. ROUTLEDGE, S.l.

48. Puljung, M., N. Vedovato, S. Usher, and F. Ashcroft. 2019. Activation mechanism of ATP- sensitive K+ channels explored with real-time nucleotide binding. eLife. 8:e41103. doi:10.7554/eLife.41103.

49. Puljung, M.C. 2021. ANAP: A versatile, fluorescent probe of ion channel gating and regulation. In Methods in Enzymology. Elsevier. 49–84.

50. R Core Team. _R: A Language and Environment for Statistical Computing_. R Core Team. _R: A Language and Environment for Statistical Computing_.

51. Resetar, A.M., and J.M. Chalovich. 1995. Adenosine 5’-(gamma-thiotriphosphate): an ATP analog that should be used with caution in muscle contraction studies. Biochemistry. 34:16039–16045. doi:10.1021/bi00049a018.

52. Sakura, H., C. Ammälä, P.A. Smith, F.M. Gribble, and F.M. Ashcroft. 1995. Cloning and functional expression of the cDNA encoding a novel ATP-sensitive potassium channel subunit expressed in pancreatic beta-cells, brain, heart and skeletal muscle. FEBS Lett. 377:338–344. doi:10.1016/0014-5793(95)01369-5.

53. Schindelin, J., I. Arganda-Carreras, E. Frise, V. Kaynig, M. Longair, T. Pietzsch, S. Preibisch, C. Rueden, S. Saalfeld, B. Schmid, J.-Y. Tinevez, D.J. White, V. Hartenstein, K. Eliceiri, P. Tomancak, and A. Cardona. 2012. Fiji: an open-source platform for biological-image analysis. Nat Methods. 9:676–682. doi:10.1038/nmeth.2019.

54. Schmied, W.H., S.J. Elsässer, C. Uttamapinant, and J.W. Chin. 2014. Efficient Multisite Unnatural Amino Acid Incorporation in Mammalian Cells via Optimized Pyrrolysyl tRNA Synthetase/tRNA Expression and Engineered eRF1. J. Am. Chem. Soc. 136:15577– 15583. doi:10.1021/ja5069728.

55. Schwanstecher, C., C. Dickel, and U. Panten. 1992. Cytosolic nucleotides enhance the tolbutamide sensitivity of the ATP-dependent K+ channel in mouse pancreatic B cells by their combined actions at inhibitory and stimulatory receptors. Mol Pharmacol. 41:480– 486.

56. Schwanstecher, C., C. Dickel, and U. Panten. 1994. Interaction of tolbutamide and cytosolic nucleotides in controlling the ATP-sensitive K+ channel in mouse beta-cells. Br J Pharmacol. 111:302–310. doi:10.1111/j.1476-5381.1994.tb14060.x.

57. Schwanstecher, C., and U. Panten. 1994. Identification of an ATP-sensitive K+ channel in spiny neurons of rat caudate nucleus. Pflugers Arch. 427:187–189. doi:10.1007/BF00585961.

58. Shandell, M.A., J.R. Quejada, M. Yazawa, V.W. Cornish, and R.S. Kass. 2019. Detection of Nav1.5 Conformational Change in Mammalian Cells Using the Noncanonical Amino Acid ANAP. Biophysical Journal. 117:1352–1363. doi:10.1016/j.bpj.2019.08.028.

59. Shyng, S.-L., and C.G. Nichols. 1997. Octameric Stoichiometry of the KATP Channel Complex. Journal of General Physiology. 110:655–664. doi:10.1085/jgp.110.6.655.

60. Sievert, C. 2020. Interactive Web-Based Data Visualization with R, plotly, and shiny.

61. Stephen, T.K.L., K.L. Guillemette, and T.K. Green. 2016. Analysis of Trinitrophenylated Adenosine and Inosine by Capillary Electrophoresis and γ-Cyclodextrin-Enhanced Fluorescence Detection. Anal Chem. 88:7777–7785. doi:10.1021/acs.analchem.6b01796.

62. Stockner, T., R. Gradisch, and L. Schmitt. 2020. The role of the degenerate nucleotide binding site in type I ABC exporters. FEBS Lett. 594:3815–3838. doi:10.1002/1873-3468.13997.

63. Treherne, J.M., and M.L. Ashford. 1992. Extracellular cations modulate the ATP sensitivity of ATP-K+ channels in rat ventromedial hypothalamic neurons. Proc Biol Sci. 247:121–124. doi:10.1098/rspb.1992.0017.

64. Tucker, S.J., F.M. Gribble, C. Zhao, S. Trapp, and F.M. Ashcroft. 1997. Truncation of Kir6.2 produces ATP-sensitive K+ channels in the absence of the sulphonylurea receptor. Nature. 387:179–183. doi:10.1038/387179a0.

65. Tusnády, G.E., É. Bakos, A. Váradi, and B. Sarkadi. 1997. Membrane topology distinguishes a subfamily of the ATP-binding cassette (ABC) transporters. FEBS Letters. 402:1–3. doi:10.1016/S0014-5793(96)01478-0.

66. Usher, S.G., F.M. Ashcroft, and M.C. Puljung. 2020a. Measuring Nucleotide Binding to Intact, Functional Membrane Proteins in Real Time. Journal of Visualized Experiments. epub ahead of print.

67. Usher, S.G., F.M. Ashcroft, and M.C. Puljung. 2020b. Nucleotide inhibition of the pancreatic ATP-sensitive K+ channel explored with patch-clamp fluorometry. eLife. 9:e52775. doi:10.7554/eLife.52775.

68. Vedovato, N., F.M. Ashcroft, and M.C. Puljung. 2015. The Nucleotide-Binding Sites of SUR1: A Mechanistic Model. Biophysical Journal. 109:2452–2460. doi:10.1016/j.bpj.2015.10.026.

69. de Wet, H., M.V. Mikhailov, C. Fotinou, M. Dreger, T.J. Craig, C. Vénien-Bryan, and F.M. Ashcroft. 2007. Studies of the ATPase activity of the ABC protein SUR1. The FEBS Journal. 274:3532–3544. doi:10.1111/j.1742-4658.2007.05879.x.

70. Wickham, H. 2016. ggplot2: Elegant Graphics for Data Analysis.

71. Zerangue, N., B. Schwappach, Y.N. Jan, and L.Y. Jan. 1999. A New ER Trafficking Signal Regulates the Subunit Stoichiometry of Plasma Membrane KATP Channels. Neuron. 22:537–548. doi:10.1016/S0896-6273(00)80708-4.

72. Zingman, L.V., A.E. Alekseev, M. Bienengraeber, D. Hodgson, A.B. Karger, P.P. Dzeja, and A. Terzic. Signaling in Channel/Enzyme Multimers: ATPase Transitions in SUR Module Gate ATP-Sensitive K؉ Conductance. 13.

